# Microbes modulate sympathetic neurons via a gut-brain circuit

**DOI:** 10.1101/545806

**Authors:** Paul A Muller, Marc Schneeberger, Zachary Kerner, Putianqi Wang, Anoj Ilanges, Fanny Matheis, Josefina del Mármol, Tiago Bruno Rezende de Castro, Matthew Perkins, Wenfei Han, Ivan de Araujo, Daniel Mucida

**Author notes:** Correspondence (P.A.M.), (D.M.). These authors contributed equally to this work (Marc Schneeberger, Zachary Kerner, Putianqi Wang).

## Abstract

Gut-brain connections monitor the intestinal tissue and its microbial and dietary content^1^, regulating both intestinal physiological functions such as nutrient absorption and motility^2-4^, and brain–wired feeding behaviour^3^. It is therefore plausible that circuits exist to detect gut microbes and relay this information to central nervous system (CNS) areas that, in turn, regulate gut physiology^5^. We characterized the influence of the microbiota on enteric–associated neurons (EAN) by combining gnotobiotic mouse models with transcriptomics, circuit–tracing methods, and functional manipulation. We found that the gut microbiome modulates gut–extrinsic sympathetic neurons; while microbiota depletion led to increased cFos expression, colonization of germ-free mice with short-chain fatty acid–producing bacteria suppressed cFos expression in the gut sympathetic ganglia. Chemogenetic manipulations, translational profiling, and anterograde tracing identified a subset of distal intestine-projecting vagal neurons positioned to play an afferent role in microbiota–mediated modulation of gut sympathetic neurons. Retrograde polysynaptic neuronal tracing from the intestinal wall identified brainstem sensory nuclei activated during microbial depletion, as well as efferent sympathetic premotor glutamatergic neurons that regulate gastrointestinal transit. These results reveal microbiota–dependent control of gut extrinsic sympathetic activation through a gut-brain circuit.

---

Extrinsic enteric–associated neurons (eEAN), comprised of sensory afferents, parasympathetic and sympathetic efferents, are equipped to sense multiple areas of the intestine simultaneously and transmit information to other tissues and to the CNS, but also complement intrinsic EANs (iEAN) in the control of gut function^6^. We sought to better characterize the connections of eEAN and whether their activity or gene expression is influenced by the gut microbiota. To identify the location of eEAN cell bodies, we injected a fluorescent retrograde tracer, cholera toxin beta subunit (CTB), into the wall of different intestinal segments, and dissected extrinsic ganglia that project to the gut, specifically the sensory nodose ganglion (NG) and dorsal root ganglia (DRG), as well as the sympathetic celiac–superior mesenteric (CG-SMG) ganglion (Fig. 1a-c, Extended Data Fig. 1a-d). Tracing specificity was confirmed by multiple approaches including subdiaphragmatic vagotomy, tracing from unrelated organs, and selective intestinal denervation (Extended Data Fig. 1e-p). Individual CTB tracing of the duodenum, ileum, and colon indicated that sensory and sympathetic innervation of these anatomically distinct intestinal regions is mediated by non-overlapping neuronal cell populations (Fig. 1d, Extended Data Fig. 1j-l). These results highlight the compartmentalization of both sensory and effector eEAN projecting to the intestine.

**Figure 1.**
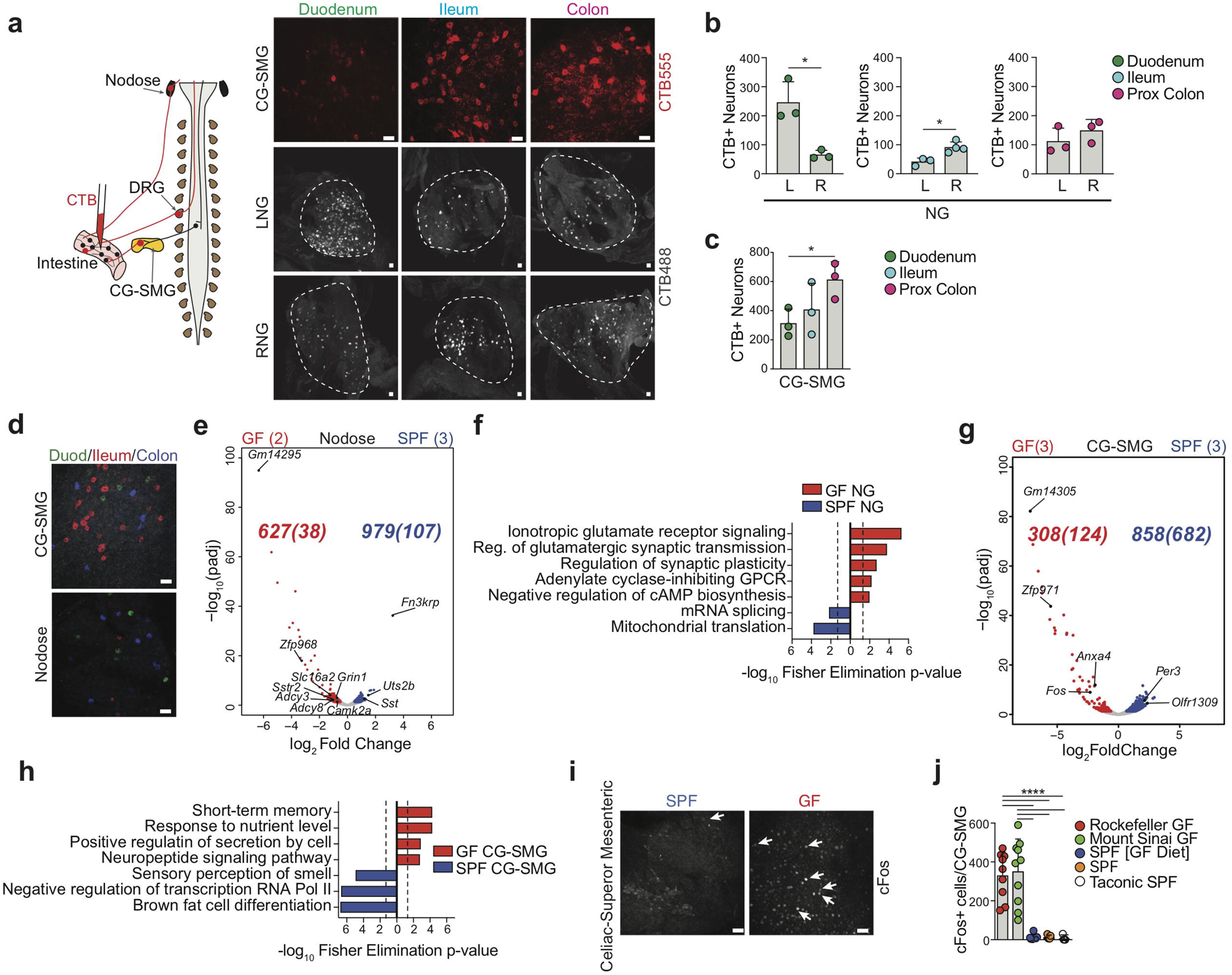
Gut sympathetic neurons are activated in the absence of a microbiota. **a**, (left) Scheme depicting and ight) representative confocal immunofluorescence images of retrograde CTB AF555 (CTB555) or AF488 (CTB488) acing from the duodenum, ileum, and colon to the CG-SMG, left (L) and right (R) nodose ganglion (NG) of 57BL/6J SPF mice. Scale bars = 50µm. Images representative of duodenum (n = 5), ileum (n = 6), and colon (n = 5). **b**, Quantification of the number of neurons in the L-NG and R-NG retrograde labeled from the duodenum, ileum, and proximal colon. * *P* < 0.05 as calculated by unpaired t-test. **c**, Number of CTB+ neurons per CG-SMG retrograde beled from the duodenum, ileum, and proximal colon.* *P* < 0.05 as calculated by unpaired t-test. **d**, Triple CTB tracing in the CG-SMG and NG with CTB488 (duodenum), CTB555 (ileum), and CTB647 (colon) from C57BL6/J SPF 1ice. Scale bars = 50µm. Images representative of n= 2. **e**, Volcano plots of differentially expressed genes compared between the NG of Snap25^RiboTag^ GF or SPF mice. Grey dots indicate all genes analyzed by differential expression analysis. Black dots indicate select genes. Red dots and number (log_2_FoldChange > 0.5) indicate genes higher in reptomycin-treated samples. Blue dots and number (log_2_ FoldChange > 0.5) indicate genes significantly higher in BS-treated samples. Parentheses indicate log_2_ Fold Change > 1 genes. Number of samples analyzed are indicated parentheses at top. **f**, Gene ontology pathways, identified by TopGO analysis of differentially expressed genes log_2_Fold Change> 0.5, padj < 0.05), enriched in the GF NG (red) vs SPF NG (blue). Dashed lines represent thresh-d of significance (1.3) as calculated by Fisher’s test with an elimination algorithm. **g**, Volcano plots of differentially expressed genes compared between the CG-SMG of Snap25^RiboTag^ GF or SPF mice. Grey dots indicate all genes 1alyzed by differential expression analysis. Black dots indicate select genes. Red dots and number (log_2_Fold-hange > 0.5) indicate genes higher in GF samples. Blue dots and number (log_2_FoldChange > 0.5) indicate genes significantly higher in SPF samples. Parentheses indicate log_2_Fold Change > 1 genes. **h**, Gene ontology pathways, identified by TopGO analysis of differentially expressed genes (log_2_Fold Change> 0.5, padj < 0.05), enriched in the F CG-SMG (red) vs SPF CG-SMG (blue). Dashed lines represent threshold of significance (1.3) as calculated by Fisher’s test with an elimination algorithm. **i**, CG-SMG whole-mount immunostaining of C57BL/6J GF and SPF mice using anti-cFos antibody. White arrows indicate cFos+ nuclei. Scale bars = 50µm. Images representative of GF (n = 9) and SPF kept in germ-free diet (n = 7). **j**, Number of cFos+ neurons in the CG-SMG of C57BL/6J GF mice from Rockefeller (n = 9) or Mount Sinai (n = 10) animal facilities as compared to SPF kept on germ-free diet (n = 7), SPF normal 10w (n = 5), and Taconic SPF normal chow (n = 5) controls. **** *P* < 0.0001 as calculated by one-way ANOVA with Jkey multiple comparisons.

We characterized microbial-mediated EAN gene expression changes by transcriptionally profiling eEAN ganglia identified by CTB-tracing using cell sorting-independent translating ribosomal affinity purification (TRAP)^7^. We interbred pan–neuronal *Snap25*^Cre^ with *Rpl22*^lsl-HA^ (RiboTag) mice^8^, which express a hemagglutinin (HA)–tagged ribosomal subunit 22, allowing for neuronal-specific immunoprecipitation of actively translated polysome-bound mRNA from whole tissue homogenates, avoiding possible confounding effects of neuronal dissociation on gene expression. Specific expression of HA–tagged ribosomes in neurons was confirmed in different eEAN ganglia isolated from *Snap25*^RiboTag^ mice (Extended Data Fig. 2a, b). We performed TRAP-seq of the NG, thoracic 9 DRG, and CG-SMG isolated from specific pathogen free (SPF) mice and *Snap25*^RiboTag^ mice re-derived into germ-free (GF) conditions (Extended Data Fig. 2c, d). Comparisons of eEAN nodes revealed significant enrichment of transcripts associated with both sensory and sympathetic neurons^9^, validating our TRAP approach (Extended Data Fig. 2c-h). We did not observe significant changes in expression of actively–translated genes in DRG between SPF and GF groups (Extended Data Fig. 2i). However, gene ontology (GO) analysis of the NG suggested an enrichment for genes associated with synaptic signalling and neuronal activation in GF mice (Fig. 1e, f). Additionally, the CG-SMG from GF animals displayed enriched GO pathways for plasticity and signalling with significantly higher transcript levels of *Fos* (Fig. 1g, h): a classic neuronal immediate–early gene and indirect marker for neuronal activity, or plasticity (used thereof interchangeably^10^). Immunofluorescence analysis confirmed that CG-SMG isolated from GF mice displayed significantly more cFos+ neuronal nuclei than their SPF mice counterparts (Fig. 1i, j, Extended Data Fig. 2j-l). Altogether these results point that absence of a microbiota results in elevated levels of gut–extrinsic sympathetic activity.

To address whether specific microbes could mediate tonic suppression of CG-SMG neurons, we utilized multiple microbial manipulation strategies. Faecal transfer from age-, sex-, strain-, and diet–matched SPF donors into GF mice restored CG-SMG neuronal cFos to levels comparable to SPF conditions, suggesting that microbiota can suppress gut–extrinsic sympathetic neurons (Fig. 2a). The mere presence of live bacteria was not enough to suppress CG-SMG neuronal activation, as evidenced by the comparable cFos levels observed in GF mice mono-colonized with segmented filamentous bacteria and GF mice (Fig. 2b). However, colonization of GF mice with the defined altered–Schaedler–flora consortium (ASF)^11^ or a defined *Clostridium spp.* consortium^12^, was sufficient to suppress cFos expression in the CG-SMG (Fig. 2b, c). Conversely, microbiota depletion of SPF mice for 2 weeks using broad-spectrum antibiotics resulted in increased cFos+ neurons in the CG-SMG when compared to control mice (Fig. 2d, e). Treatment with either ampicillin or vancomycin, but not with metronidazole or neomycin, was sufficient to drive sympathetic cFos expression, suggesting that specific subsets of bacteria were able to suppress cFos activation (Fig. 2f). Additionally, a single oral gavage of streptomycin to SPF C57BL/6, BALB/c, or CBA/J mice resulted in CG-SMG neuronal activation as early as 24 h post-gavage, as well as decrease in microbiota diversity. Five days post streptomycin treatment, cFos levels in the CG-SMG returned to SPF levels, indicating that gut–sympathetic activity is readily tuned to changes in the gut microbiota (Fig. 2g, Extended Data Fig. 2m, n). Finally, to address whether activated CG-SMG sympathetic neurons project to the intestine, we injected fluorescent CTB in the ileum of broad-spectrum antibiotic–treated *Fos*^GFP^ mice. We observed extensive colocalization between CTB+ (red) and cFos+ (green) neurons in the CG-SMG (Fig. 2h), confirming that sympathetic neurons activated upon microbial depletion project to the intestine, although we cannot exclude projections to other visceral tissue connected to the CG-SMG. The above results indicate that a defined microbial consortium is sufficient to suppress cFos expression in gut sympathetic neurons, and that sympathetic activity can reflect shifts in the gut microbial community.

**Figure 2.**
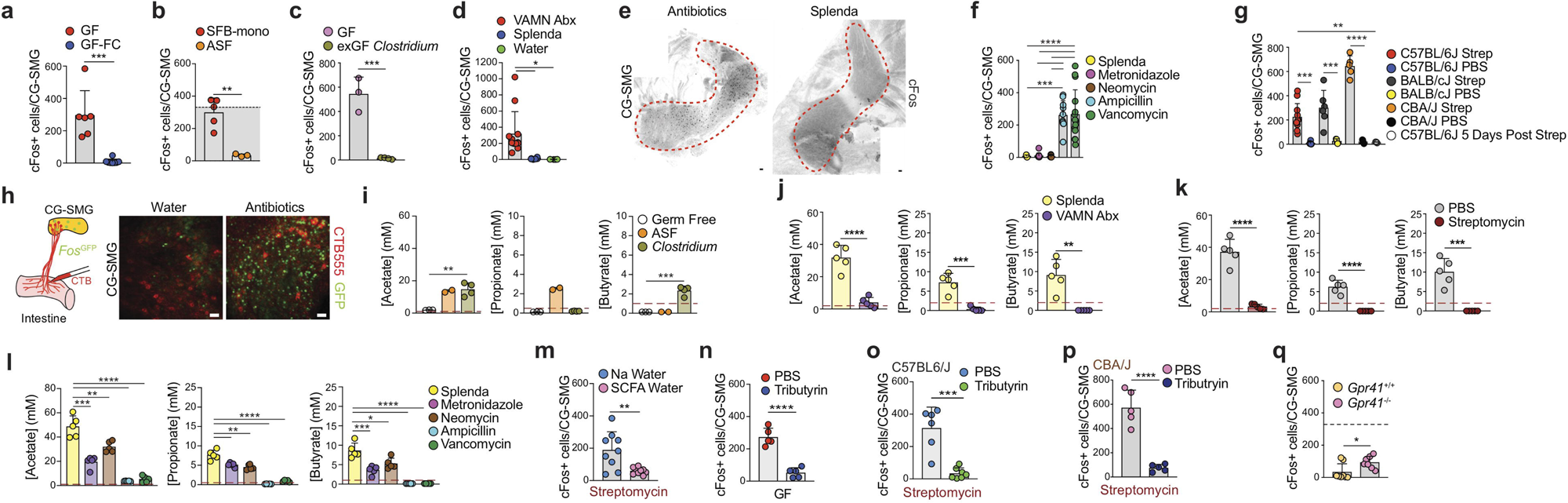
Microbe-derived signals suppress gut sympathetic neurons. **a**, Number cFos+ neurons in the CG-SMG of C57BL/6J GF mice (n = 6) and GF mice (n = 8) colonized ith feces from SPF mice 2 weeks prior to analysis. *** *P* < 0.001 as calculated by unpaired t-test. **b**, Number of cFos+ neurons in the CG-SMG of C57BL/6J SFB monocolonized ice (n = 5) and ASF colonized mice (n = 3). Dashed line and shaded area below indicate the average number (334) of cFos+ neurons in GF mice. ** *P* < 0.01 as calculated by 1paired t-test. **c**, Number of cFos+ neurons in the CG-SMG of Germ Free C57BL/6J mice from Weill Cornell with (n = 4) or without (n = 3) *Clostridium* colonization.*** *P* < 0.001 as measured by unpaired t-test. d, Number of cFos+ neurons in the CG-SMG of C57BL/6J SPF mice treated for 2 weeks with vancomycin, ampicillin, metronidazole, and neomy-n (n = 12) in drinking water, Splenda (n = 6) or water (n = 5) controls.* *P* < .05 as calculated by one way ANOVA with Tukey’s multiple comparisons. e, Whole-mount stitched images ‘cFos staining in the CG-SMG from of C57BL/6J SPF mice treated with VAMN or Splenda. Images representative of at least n= 6. Scale bars = 50µm. **f**, Number of cFos+ neurons in the CG-SMG of C57BL/6J SPF mice treated for 2 weeks with single antibiotic (metronidazole: n= 10, neomycin: n= 10, ampicillin: n= 10, vancomycin: n= 15) or Splenda (n 5). *** *P* < 0.001, **** *P* < 0.0001 as calculated by one way ANOVA with Tukey’s multiple comparisons. **g**, cFos+ neurons in the CG-SMG of C57BL/6J (streptomycin: n= 11, BS: n= 6), BALBc/J (streptomycin: n = 6, PBS: n = 4), and CBNJ (streptomycin: n = 5, PBS: n = 5) mice 24 hours post oral gavage of streptomycin or PBS and 5 days post streprnycin in C57BL/6J mice (n = 4). ** *P* < .01, *** *P* < .001, **** *P* < 0.0001 as calculated by one way ANOVA with Tukey’s multiple comparisons. **h**, Whole-mount of CG-SMG of *Fos*^GFP^ SPF mice treated with VAMN or water for 2 weeks, injected with AlexaFluor555 conjugated CTB into the ileum. Scale bars = 50µm**. i**, Quantification of SCFA in the cecal mtents of Germ Free mice colonized with ASF or *Clostridium* consortium. Data shown as mean+/- s.d. ** *P* < 0.01, *** *P* < 0.001 as calculated by one way ANOVA with Tukey’s multiple comparisons. Dashed line indicates lowest limit of detection. **j**, Quantification of SCFA in the cecal contents of VAMN- and Splenda-treated C57BL6/J mice. Data shown as mean +/- s.d. ** *P* < 0.01, *** *P* < 0.001, **** *P* < 0.001 as calculated by unpaired t-test. Dashed line indicates lowest limit of detection. **k**, Quantification of SCFA in the cecal mtents of PBS- and streptomycin-treated C57BL6/J mice. Data shown as mean +/- s.d. *** *P* < 0.001, **** *P* < 0.0001 as calculated by unpaired t-test. Dashed line indicates west limit of detection. I, Quantification of SCFA in the cecal contents of single antibiotic- and Splenda-treated C57BL6/J mice. Data shown as mean +/- s.d. * *P* < 0.05, ** *P* < 01, *** *P* < 0.001, **** *P* < 0.0001 as calculated by one way ANOVA with Tukey’s multiple comparisons. Dashed line indicates lowest limit of detection. **m**, Number of cFos+ neurons in the CG-SMG of C57BL/6J SPF mice 24 hours post oral gavage with streptomycin in mice treated with SCFA-supplemented or sodium-containing water. ** *P* < 0.01, as 3lculated by unpaired t-test. n, Number of cFos+ neurons in CG-SMG of C57BL/6J GF mice 24 hours post-treatment with tributyrin (n = 5) or PBS (n = 5). Data shown as mean +/- s.d. **** *P* < 0.0001 as calculated by unpaired t-test. **o**, Number of cFos+ neurons in CG-SMG of C57BL/6J SPF mice treated with tributyrin (n = 7) or PBS (n = 6) 24 hours post oral gavage of stremptmycin. Data shown as mean +/- s.d. *** *P* < 0.001 as calculated by unpaired t-test. **p**, Number of cFos+ neurons in CG-SMG of CBNJ SPF mice treated ith tributyrin (n = 5) or PBS (n = 5) 24 hours post oral gavage of stremptomycin. **** *P* < 0.0001 as calculated by unpaired t-test. **q**, Number of cFos+ neurons per CG-SMG in Gnr41^-/-^ mice Dashed line indicates average number of cFos+ neurons for GE mice (334) Data shown as mean +/- s.d \**P<* 0.05 calculated by unnaried t-test

We observed that gnotobiotic manipulations resulted in suppression of CG-SMG neurons when defined microbial consortia, known to restore levels of SCFAs, were introduced^11-13^. Mass spectrometric quantification of SCFAs confirmed that GF mice lacked detectable levels of butyrate, propionate, and acetate in their cecal contents (Fig. 2i). GF mice colonized with ASF displayed a significant increase in the levels of propionate and acetate in their ceca, while the *Clostridium spp.* consortium led to a significant boost in butyrate and acetate levels (Fig. 2i). Furthermore, treatment with broad-spectrum antibiotics, streptomycin, ampicillin or vancomycin alone recapitulated SCFA depletion to below detection threshold (Fig. 2j-l). In each of these cases, the presence or absence of luminal SCFAs correlated with the number of cFos+ neurons in the CG-SMG. In contrast, in the context of *Salmonella typhimurium* infection, which we previously reported to induce CG-SMG activation^14^, we did not detect significant changes in SCFAs at early stages of infection^15^, suggesting that additional signals contribute to CG-SMG activation upon infection or in inflammatory contexts (Extended Data Fig. 3a-c). Next, we tested whether supplementation of SCFAs in microbiota–depleted mice restores cFos levels in their CG-SMG. Administration of exogenous butyrate, acetate, and propionate in the drinking water suppressed streptomycin–induced cFos (Fig 2m). Moreover, administration of tributyrin, a butyrate pro-drug containing three acetylated butyrate molecules that are readily hydrolysed in the intestine^15,16^, was sufficient to suppress cFos expression in CG-SMG neurons of both GF mice and SPF mice treated with streptomycin (Fig. 2n-p, Extended Data Fig. 3d). Butyrate, and additional SCFAs, can modulate gene expression in target cells via activation of G protein-coupled receptors GPR41, 43 or 109A, inhibition of histone deacetylases or act as an energy substrate^17^. We evaluated cFos+ neurons in the CG-SMG in mice deficient for GPR41, GPR109A or double-deficient for GPR43 and GPR109A, all maintained under SPF conditions. While Gpr109a^−/−^ and double Gpr43^−/−^ Gpr109a^−/−^ mice did not show differences when compared to wild-type controls, Gpr41^−/−^ mice showed a mild yet significant increase in the number of cFos^+^ neurons in the CG-SMG (Fig. 2q, Extended Data Fig. 3e, f). These data suggest a potential role for signalling via GPR41, which is expressed by intestinal epithelial cells, iEAN, and eEAN^18^, as an upstream pathway modulating gut sympathetic ganglia. Investigation of additional microbe-modulated epithelial cell factors revealed that GLP-1^16^ and PYY^19^ could modulate sympathetic activity independently, or in the context of microbiota depletion, respectively (Supplementary Information 1, Extended Data Fig. 4a-i). Overall, the above results identify SCFAs as physiological modulators of gut sympathetic neuronal activation.

We investigated the neuronal population(s), or circuits, upstream of the CG-SMG that could be involved in driving sympathetic activity under conditions of microbial depletion. Our analyses did not support a role for direct sensing of microbial depletion by CG-SMG sympathetic neurons, nor for viscerofugal neurons modulating gut sympathetic activity (Supplementary Information 2, Extended Data Fig. 5a-t, Supplementary Video 1-2). Thus, we investigated whether premotor neurons in brain areas could receive sensory input originating in the intestine to then modulate the sympathetic output. We focused on defining whether sympathetic premotor areas in the brainstem could modulate the CG-SMG through sympathetic preganglionic neurons in the spinal cord (SPN)^20^. To define these premotor areas, we performed polysynaptic retrograde tracing by injecting mRFP1-expressing pseudorabies virus PRV-614 ^21^ into the ileum or colon of SPF mice. We confirmed that viral spread moved back from the intestine to CG-SMG, then to SPN in the spinal cord (Fig. 3a-d, Supplementary Video 3-5)^22^. Analysis of the brains at day 4 post-injection of PRV-RFP by AdipoClear tissue clearing (using a modification of Clearmap analysis^23^) identified the caudal raphe nuclei, involved in thermogenesis and various gastrointestinal functions^24,25^; the gigantocellular reticular nucleus (Gi) and the lateral paragigantocellular nucleus (LPGi), both thought to be involved in locomotion and gastrointestinal function^22,25^, and the rostroventrolateral medulla (RVLM), involved in autonomic regulation via C1 neurons^26^ (Fig. 3e, f, Extended Data Figure 6a, Supplementary Video 6). The specificity of this polysynaptic tracing was verified by comparison to intraperitoneal PRV injection or ileal PRV injection coupled with subdiaphragmatic vagotomy (sdVx), which eliminates brainstem areas labelled via vagal motor neurons (Extended Data Fig. 6b-d). Search of the Allen Brain Atlas *in situ* hybridization database for potential neurotransmitter-defined populations in these sympathetic premotor areas pointed to inhibitory GABAergic neurons and excitatory glutamatergic neurons as possible populations involved in the modulation of downstream circuits polysynaptically connected to the intestine. Injection of PRV-RFP into the ileum of *Slc32a1*^L10-GFP^ (VGAT: inhibitory) and *Slc17a6*^L10-GFP^ (VGLUT2: excitatory) mice revealed extensive colocalization of RFP+/GFP+ neurons, with a majority of Gi neurons identified as VGAT+, and LPGi/RVLM neurons identified as VGLUT2+ respectively with a minor contribution from other neurotransmitters (Fig. 3g-j, Extended Data Fig. 6e. To address whether signals from different intestinal regions diverge from distinct or common neuronal populations in the brain, we injected PRV-RFP into the ileum and PRV-GFP into the colon or duodenum of wild-type SPF mice. A majority of neurons in each of these four brainstem areas were labelled with both fluorescent proteins (Fig. 3k, Extended Data Fig. 6f). These experiments define efferent premotor brainstem neurons polysynaptically connected to different regions of the intestine via preganglionic neurons in the spinal cord that, in turn, may control gut–sympathetic activity.

**Figure 3.**
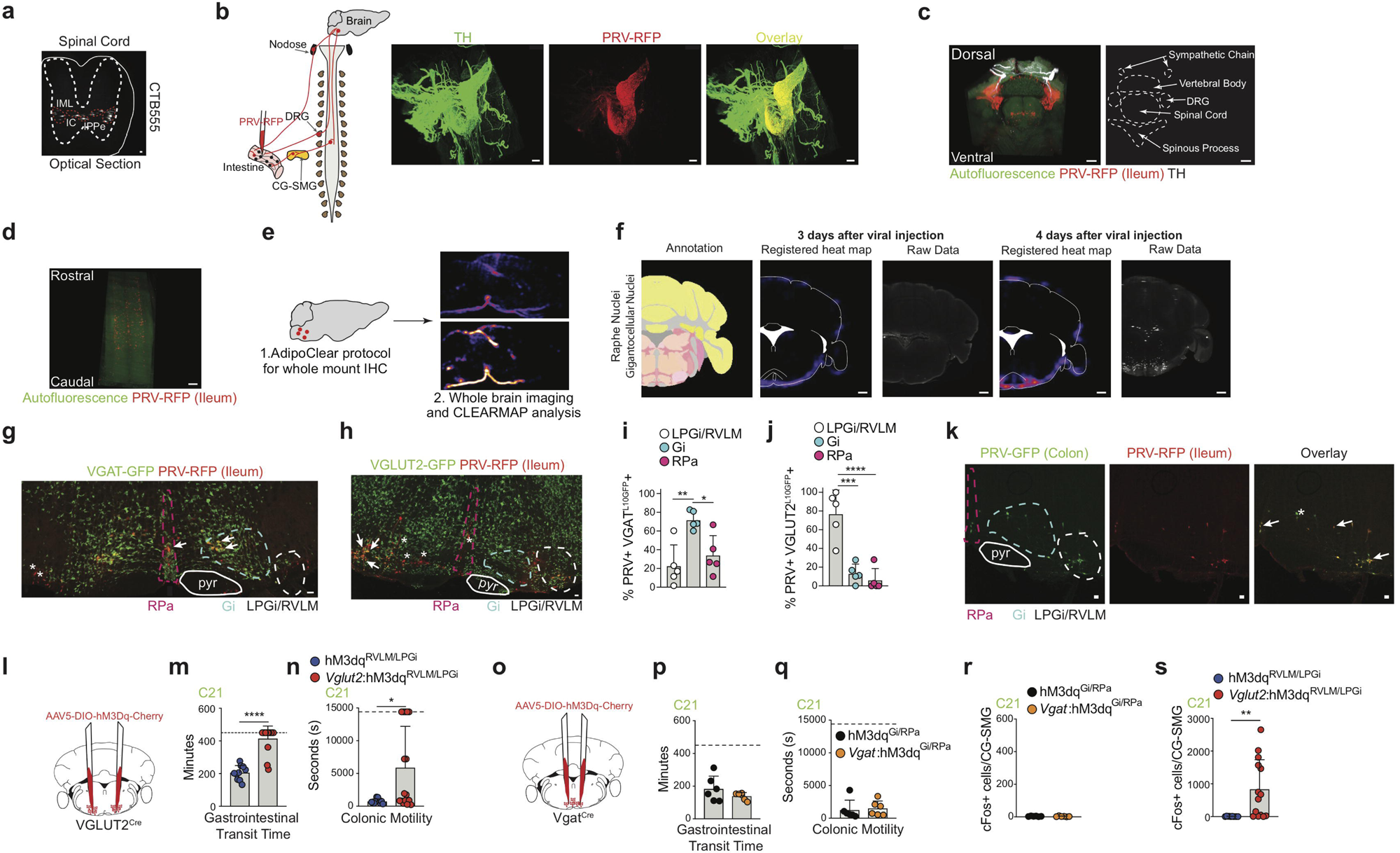
The efferent arm of microbe-modulated gut sympathetic neuronal circuit. **a**, Optical projection of 3DISCO-cleared spinal cord showing sympathetic preganglionic neurons retrograde labeled by injection of CTB555 into the CG-SMG of C57BL/6J SPF mice. Scale bar = 20µm. IML= intermediolateral cell column, IC: intercalated nucleus, IPPe: central autonomic area. Image representative of n= 3. **b**, (left) Scheme representing injection of PRV into the intestine for 1ultisynaptic tracing (for b-1). (right) 3D reconstruction of calAdipoClear-cleared CG-SMG with PRV-RFP (red) injected into the ileum 4 days post injection, stained with anti-TH (green) antibody. Scale bar = 300µm. Image representative of n= 3. **c**, (left) 3D reconstruction of calAdipoClear-cleared intact spinal column ith PRV-RFP (red) inected into the ileum 4 days post injection, stained with anti-TH (white) antibody. Autofluorescence in green. (right) outline of relevant spinal xd structures. Scale bars = 400µm. Image representative of n= 3. **d**, 3D-cropped image of the spinal cord from c, showing sympathetic preganglionic neurons beled by PRV-RFP injected into the intestine. Scale bar = 300µm. Image representative of n= 3. **e**, (right) Scheme representing tissue clearing followed by learMap analysis. f, ClearMap analysis results of PRV-RFP injection into the ileum at days 3 and 4 post-injection showing raphe/gigantocellular nuclei. Scale 3r = 1mm. Images representative of n= 3. **g**, Confocal image of a vibratome section of the brainstem showing native VGAT^L10GFP^ and PRV-RFP fluorescence 4 3ys post PRV injection. Relevant brainstem regions are highlighted. Arrows indicate GFP+ RFP+ cells and asterisks indicate RFP+ GFP-cells. Scale bar = 50 µm. Image representative of n = 5. **h**, Confocal image of a vibratome section of the brainstem stained with anti-GFP (VGLUT2 ^L10GFP^) and anti-RFP (PRV-RFP) 1tibodies 4 days post-injection of PRV. Relevant brainstem regions are highlighted. Arrows indicate GFP+ RFP+ cells and asterisks indicate RFP+ GFP-cells. cale bar = 50µm. Image representative of n= 5. i, Percentage of PRV+ VGAT^L10GFP^ + neurons in RPa, Gi, and LPGi/RVLM 4 days post injection. (n = 5). Data 1own as mean+/- s.d. * *P* < 0.05, ** *P* < 0.01 as calculated by one way ANOVA with Tukey’s multiple comparisons. **j**, Percentage of PRV+ VGLUT2 ^L10GFP^+ neurons in RPa, Gi, and LPGi/RVLM 4 days post injection. (n = 5). Data shown as mean+/- s.d. *** *P* < 0.001, **** *P* < 0.0001 as calculated by one way ANOVA with ukey’s multiple comparisons. **k**, Confocal image of a vibratome section of the brainstem showing PRV-GFP (duodenum) and PRV-RFP (ileum) 4 days post-in- ction. Arrows indicate GFP+ RFP+ cells and asterisks indicate RFP+ GFP- cells. Images representative of n= 3. Scale bars = 50µm. pyr = pyramidal tract, RPa raphe pallidus, Gi = gigantocellular nucleus, LPGi = lateral paragigantocellular nucleus, RVLM = rostral ventrolateral medulla. **l**,**o**, Scheme representing the jection of AAV5-DIO-hSyn1-hM3Dq-mCherry into the LPGi/RVLM of VGLUT2^cre^ mice (I) or into the Gi/RPa of Vgat^cre^ mice **(o). m**,**n**,**p-s**, Gastrointestinal transit time **(m**,**p)**, colonic motility **(n**,**q)** and number of cFos+ neurons in the CG-SMG **(r**,**s)** of Vgat:hM3Dq ^Gi/RPa^and VGLUT2:hM3Dq ^LPGi/RVLM^mice treated with 1mg/kg Compound 21. Dashed line indicates maximum time allowed per animal for motility measurement. * *P* < .05, ** < .01, **** < .0001 as calculated by unpaired t-test

Gut sympathetic innervation is involved in the control of gut motility and secretion^20,25^. Consistent with a possible role for the premotor brainstem nuclei identified above in the regulation of gut sympathetic activity, we observed elevated cFos in both RPa and LPGi/RVLM neurons in GF mice (Extended Data Fig. 7a), which are known to display gut dysmotility^27^. To identify and permanently label recently-activated neurons following antibiotic treatment, we combined *Fos*^TRAP2:tdTomato^ mouse plasticity mapping^28^ and fluorescent PRV injection into the proximal colon, labelling gut–projecting neurons. Consistent with the increase in cFos, we observed an increase in the percentage of gut–connected (PRV+) TRAP+ cells in the LPGi/RVLM of these mice (Extended Data Fig. 7b). To directly determine whether these brainstem populations modulate gut–sympathetic activity, we bilaterally injected excitatory AAV5-DIO-hSyn-hM3Dq-mCherry into the Gi or the LPGi/RVLM of VGAT^Cre^ and VGLUT2^Cre^ mice, respectively (Extended Data Fig. 7c). Designer Receptors Exclusively Activated by Designer Drugs (DREADD) enable sustained chemogenetic modulation of neurons using the specific designer ligand compound 21 (C21)^29^. C21 administration to wild-type or (Gi)^VGAT^ mice did not affect baseline motility measurements, but it significantly slowed intestinal transit and faecal pellet output in (LPGi/RVLM)^VGLUT2^ mice (Fig. 3l-q, Extended Data Fig. 7d-f). As an additional control for the accurate targeting of these brain regions^22^, C21 administration affected locomotion in both VGLUT2 and VGAT groups (Extended Data Fig. 7g, h). Finally, consistent with a role for the excitatory nuclei upstream of gut-sympathetic activation, chemogenetic activation of (LPGi/RVLM)^VGLUT2^ neurons, but not of (Gi)^VGAT^, led to a significant increase in the number of cFos+ neurons in the CG-SMG (Fig. 3r, s). These findings demonstrate that glutamatergic LPGi/RVLM brainstem neurons are capable of driving gut sympathetic activity, which in turn can slow gastrointestinal transit.

We investigated further the ClearMap PRV-RFP analysis described above to identify additional brain areas synaptically connected to the gut, and possibly to the LPGi/RVLM. Examination of data four days post PRV infection revealed additional brainstem regions previously shown to connect to the stomach^30^ and rectum^31^. Specifically, we observed significant PRV labelling in the dorso-vagal complex^31^ including the dorsal motor nucleus of the vagus (DMV), nucleus tractus solitarius (NTS) and area postrema (AP) (Extended Data Fig. 8a-d). Previous studies demonstrated that the NTS and AP can directly integrate gut sensory information from vagal sensory neurons or circulating factors^3,32^ and that these nuclei connect to the LPGi/RVLM^33,34^. To determine whether changes in microbial load could affect neuronal activity in the NTS and AP, we measured cFos in these areas after antibiotic treatment. Following streptomycin treatment, we observed a significant increase in cFos expression in both the NTS and AP (Fig. 4a, Extended Data Fig. 8e). These areas also displayed high levels of cFos in GF mice and *Fos*^TRAP2:tdTomato^ treated with streptomycin, further suggesting a functional relevance of these areas (Extended Data Fig. 8f-h). To assess whether these sensory brainstem nuclei are involved in the detection of SCFAs, we analysed their cFos levels post administration of tributyrin to streptomycin–treated mice. While cFos levels in the AP were suppressed, levels in the NTS remained elevated (Fig. 4b, Extended Data Fig. 8i), suggesting the AP as a distal sensory hub for intestinal SCFAs, although additional SCFA and unknown visceral signals might be sensed in the NTS during dysbiosis or microbial depletion. These results characterize a set of brainstem sensory nuclei tuned to detect changes in the gut microbiota or metabolites thereof.

**Figure 4.**
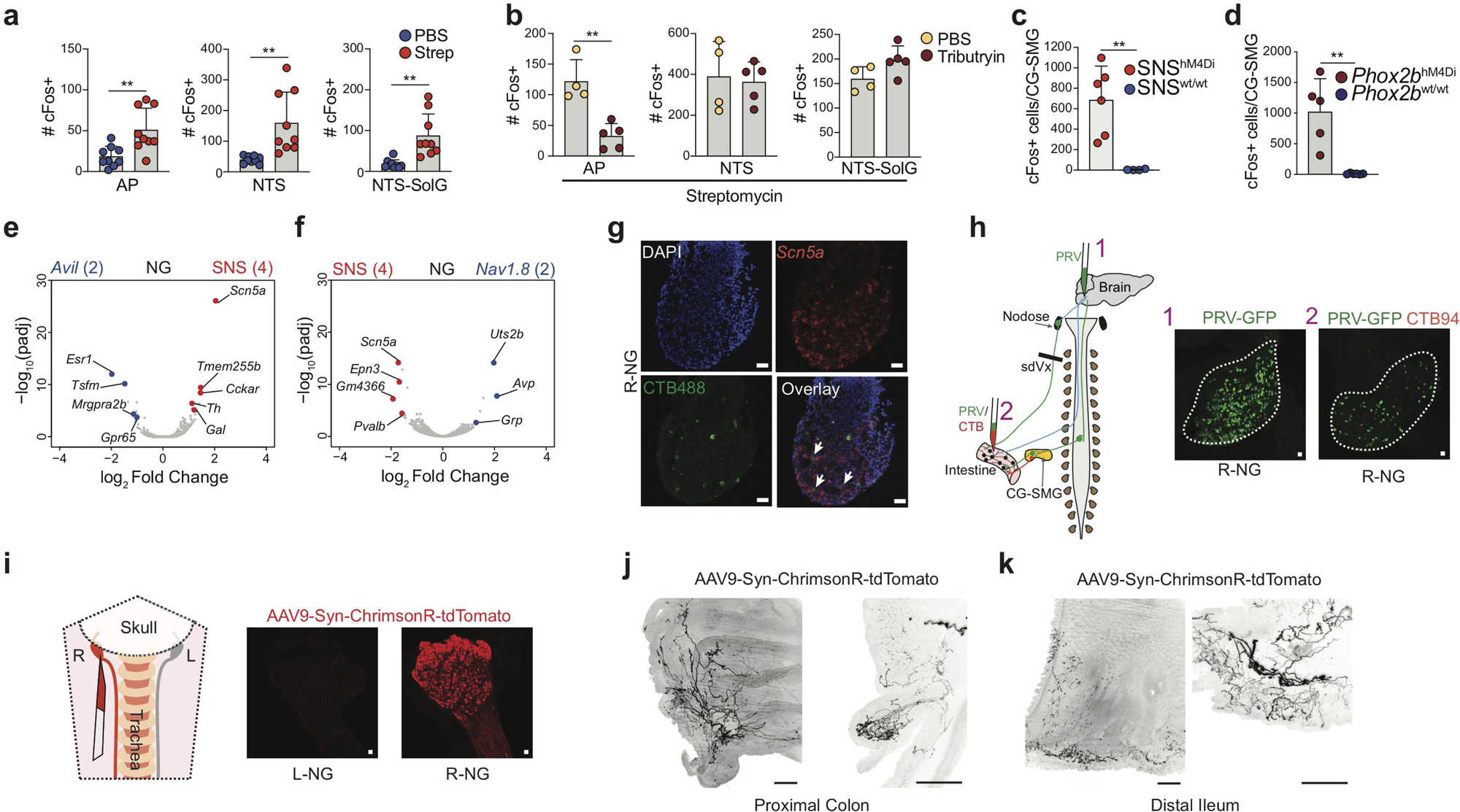
The afferent arm of microbe-modulated gut sympathetic neuronal circuit. **a**, Number of cFos+ neurons in the AP, TS-gelatinous (SolG), and NTS inclusive of SolG, of C57BL/6J SPF mice 24 hours post oral gavage with PBS or streptomycin.\*\**P* < 0.01, as calculated by unpaired t-test. **b**, Number of cFos+ neurons in the AP, NTS, and SolG of C57BL/6J SPF mice 24 hours post oral gavage of streptomycin followed by treatment with PBS or Tributyrin. ** *P* < 0.01, as calculated by unpaired t-test. **c**, Number of cFos+ neurons in the CG-SMG of SNS ^hM4Di^ (n = 6) or control mice (n = 4) 6 hours post 10mg/kg C21 injection. Data 1own as mean +/- s.d. ** *P* < 0.01 as calculated by unpaired t-test. **d**, Number of cFos+ neurons in the CG-SMG of Phox2b^hM4Di^ 1= 5) and *Phox2b*^wt/wt^ mice (n = 6) 4 hours post 10mg/kg C21 injection. Data shown as mean +/- s.d. ** P < 0.01 as calculated by 1paired t-test. **e**, Volcano plot of differentially expressed genes from the nodose ganglion of SNS^RiboTa9^ and A*vil*^RiboTag^ mice. Blue dots indicate genes significantly enriched in the NG of A*vil*^Ribo Tag^ mice. Red dots indicate genes significantly enriched in the NG of NS^Ribo Tag^ mice. Number of samples analyzed are indicated in parentheses at top. **f**, Volcano plot of differentially expressed genes Jm the NG of SNS ^Ribo Tag^and Nav1.8^RiboTag^ mice. Blue dots indicate genes significantly enriched in the NG of Nav1.8^Ribo Tag^ mice. Red dots indicate genes significantly enriched in the NG of SNS ^Ribo Tag^ mice. Number of samples analyzed are indicated in parentheses at top. **g**, RNAscope *in situ* hybridization images of the R-NG from C57BL/6J SPF mice injected with CTB488 into the proximal coIon, showing *Scn5a (Nav1.5)* (red) and immunofluorescence for CTB488. White arrows point to CTB+ *Scn5a+* cells. Images representative of n= 2. Scale bars = 50µm. **h**, Scheme of PRV (1) or PRV and CTB co-injection (2). Whole-mount IF image of the G from C57BL/6J SPF injected with PRV-GFP in the brainstem (left, 1), or injected along with CTB94 in the proximal colon in mice with sdVx (right, 2). Scale bars = 50μm. Outline hightlights NG. Scale bars = 50μm. **i**, (left) Scheme of right NG injection. (right) Expression of AAV9-Syn-ChrimsonR-tdTomato in NG. Scale bars = 50μm. j, **k**, Whole-mount cleared-tissue IF image of the proximal colon (**j**) or the distal ileum (**k**) of mice injected with AAV9-Syn-ChrimsonR-tdTomato in the R-NG (left 1x and right 4x zoom). Scale bars = 500μm (left) and 250μm (right). NG = nodose ganglion, RPa = raphe pallidus, Gi = gigantocellular nucleus, LPGi = lateral paragigantocellular nucleus, RVLM = rostral ventrolateral medulla, AP = area postrema, NTS = nucleus tractus soli-tarius.

The increased cFos expression in the NTS/AP upon microbial depletion suggests a functional role for afferent sensory neurons in the regulation of gut sympathetic activity. Extrinsic neuronal sensing at the intestinal epithelium, mucosa, and *muscularis* is primarily carried out by sensory afferents in the NG and DRG^35^, both also identified by our retrograde strategies described above. To target NG or DRG sensory activity we first crossed SNS^Cre^, a BAC-transgenic mouse expressing Cre under the *Scn10a* (*Nav1.8*) promoter^36^, with inhibitory hM4Di (*Rosa26*^lsl-hM4Di^) mice. Therefore if afferent eEAN were involved in this circuit, administration of C21 to SNS^hM4Di^ mice should result in the inhibition of afferent eEAN^37^, functionally phenocopying the loss of microbial signals and thus resulting in activation of gut sympathetic neuron activation. We confirmed that SNS^Cre^ recombination targets most DRG and NG neurons, in addition to robust labelling of nerve fibres spanning the width of the intestinal wall (Extended Data Fig. 9a, b, Supplemental Table 1). Indeed, SNS^hM4Di^ mice displayed a significant activation of CG-SMG neurons upon administration of C21 (Fig. 4c). To determine whether NG alone or combined NG/DRG inhibition was responsible for this phenotype, we crossed *Rosa26*^lsl-hM4Di^ with *Phox2b*^Cre^ mice, a Cre line that specifically targets the NG, and a small population of iEAN, while avoiding recombination of DRG neurons^38,39^ (Extended Data Fig 9a, b, Supplemental Table 1). Injection of C21 into *Phox2b*^hM4Di^ mice also led to a marked increase in the number of cFos+ cells in the CG-SMG (Fig. 4d). Because both SNS^hM4Di^ and *Phox2b*^hM4Di^ also target a population of CG-SMG neurons, we performed selective chemogenetic manipulation of CG-SMG neurons directly, which ruled out possible hM4Di-mediated activation (Supplementary Information 3, Extended Data Fig. 9c-f). These chemogenetic experiments indicate that modulation of vagal afferents is sufficient to drive gut sympathetic activity.

To narrow down the potential population of NG neurons targeted by these approaches, and their role in this gut-brain-gut loop, we first isolated the NG from SNS^RiboTag^ mice, the Cre line in which DREADD modulation resulted in CG-SMG activation, or from *Avil*^RiboTag^ and *Nav1.8*^RiboTag^ mice, i.e. Cre lines that labelled subsets of NG neurons but were not affected by DREADD modulation (Extended Data Fig. 9g-i). TRAP-seq analysis revealed a significant enrichment in the gene *Scn5a* (*Nav1.5*) in the NG of SNS^RiboTag^ mice when compared to either *Avil*^RiboTag^ or *Nav1.8*^RiboTag^ mice (Fig. 4e, f). Moreover, *in situ* hybridization coupled with intestinal retrograde tracing confirmed that a proportion of gut-projecting NG are indeed Nav1.5+ and targeted by SNS^Cre^ mice (Fig. 4g, Extended Data Fig. 9j-l). To address whether vagal modulation can activate the sensory brainstem nuclei that we found to be modulated by microbiota manipulation, we examined the NTS and AP of SNS^hM4Di^ and *Phox2b*^hM4Di^ mice treated with C21, and observed significant increase in cFos (Extended Data Fig. 10a, b). We chose this indirect measure of vagal activity as reported markers of intrinsic NG activation in mice, such as phosphorylated ERK^40,41^, are not well characterized and vagal nerve recordings lack the resolution required to detect physiologic changes in afferent activity. We were not able to determine whether this increase in cFos is the result of hM4Di activation or simply disinhibition of NTS/AP neurons. Additionally, we were unable to surgically assess the requirement of vagal sensory input to these areas, as sdVx alone led to a significant increase in NTS/AP cFos (Extended Data Fig. 10c, d). However, bilateral injection of fluorescent PRV into the LPGi/RVLM confirmed that both connect to the NG (Fig. 4h, Extended Data Fig. 10e). Conversely, injection of fluorescent PRV into the wall of the colon of sdVx mice resulted in significant labelling of the left and right NG, with a bias towards right NG labelling, mirroring the trend observed with CTB tracing from the distal intestine. These results confirm the identification of a gut-vagal afferent sensing circuit that can potentially drive sympathetic activity and is distinct from the vagal parasympathetic circuitry (Fig. 4h, Extended Data Fig. 10f, g). Finally, visualization of vagal afferents projections using Cre–dependent (SNS^Cre^ and VGlut2^Cre^) and -independent (direct injection) AAV anterograde tracing methods^42-44^, revealed dense labelling of neuronal processes particularly in the distal intestine, suggesting that NG sensory afferent neurons project fibres to sites of high microbial load or SCFA concentration (Fig. 4i-k, Extended Data Fig. 10h-j and Supplementary Video 7-11). Overall, our results identify a gut-brain-gut circuit whereby distinct microbes and microbial metabolites modulate activation of gut sympathetic neurons and brainstem sensory nuclei capable of integrating gut-specific stimuli (Extended Data Fig. 10k).

The functional, circuit and gene expression–based studies presented here suggest that EAN detection of microbes or their metabolites, either directly or indirectly via epithelial cells, is a core sensory system whereby alterations in microbial composition is sufficient to significantly activate gut–projecting neurons. Gut sympathetic activity can impact blood flow, motility, and epithelial secretion^25,45^, which in turn may influence microbial niches in the lumen. Activation of the gut-sympathetic nervous system following microbial depletion may also explain the anti-inflammatory effects of specific types of antibiotics^46^. Additional CG-SMG targets such as the spleen, pancreas, and liver may also be regulated by the microbiota with conceivable impacts systemic immunity and metabolism. Furthermore, sympathetic signalling can impact gene transcription in a variety of cell targets found in the intestine and elsewhere, including gut–resident macrophages and innate lymphoid cells^14,47^. Our data also suggests that a particular population of gut–projecting vagal sensory neurons, enriched in transcript for the voltage gated sodium channel *Nav1.5*, may play a role in mediating CNS–driven sympathetic suppression. It is intriguing to note that mutations in *Nav1.5* have been described as a hallmark of a subset of irritable bowel syndrome (IBS) patients with constipation^48,49^. It is therefore plausible that for specific IBS cohorts, dysbiosis and changes in the local availability of SCFAs or additional microbial metabolites result in increased sympathetic activity, driving slower intestinal motility. We identified multiple potential microbiota– derived signals that can modulate gut sympathetic activity and neuronal populations synaptically connected to the gut. In light of our findings, further characterization of microbial regulation of the autonomic nervous system, and additional circuits that integrate microbial signals, may be key for understanding the regulation of intestinal motility, visceral pain, enteric immunity, and systemic disorders related to the gut–brain axis; knowledge essential for defining therapeutic strategies.

## Methods

### Mice

Mice were housed in a 12hr light-dark cycle with *ad libitum* access to food and water. Wild-type mice used: C57BL/6 (C57BL/6J, Jackson #000664 or C57BL/6NTac, Taconic #B6-M/F), CBA/J (Jackson #000656), BALB/cJ (Jackson #000651). Transgenic mice used: *Fos*^GFP^ (B6.Cg-Tg(Fos/EGFP)1-3Brth, Jackson #014135), RiboTag (B6N.129-*Rpl22*^*tm1.1Psam*^, Jackson #011029), *Snap25*^cre^ (B6;129S-*Snap25*^*tm2.1(cre)Hze*^, Jackson #023525), *Rosa26*^lsl-hM4Di^ (B6N.129-*Gt(ROSA)26Sor*^*tm1(CAG-CHRM4*,-mCitrine)Ute*^, Jackson #026219), *Rosa26*^lsl-tdTomato^ (B6.Cg-*Gt(ROSA)26Sor*^*tm14(CAG-tdTomato)Hze*^, Jackson #007914), *Vgat*^cre^ (*Slc32a1*^*tm2(cre)Lowl*^, Jackson #016962), *Chat*^cre^ (B6;129S6-*Chat*^*tm2(cre)Lowl*^, Jackson #006410). SNS^cre^ (Tg(*Scn10a*^cre^)1Rkun, gift of R. Kuhner), *Nav1.8*^cre^ (*Scn10a*^tm2(cre)Jnw^, gift of J. Wood), *Vgat*^cre^ (*Slc32a1*^*tm2(cre)Lowl*^, Jackson #016962), *Advillin*^CreERT2^ (B6.Cg-Tg(*Avil*-icre/ERT2)AJwo/J Jackon # 026516), *Vglut2*^cre^ (*Slc17a6*^*tm2(cre)Lowl*^/J, Jackson #016963), *Glp1r*^cre^ (*Glp1r*^*tm1.1(cre)Lbrl*^/J, Jackson #029283), *Phox2b*^cre^ (B6(Cg)-Tg(Phox2b-cre)3Jke/J, Jackson #016223), *Villin*^creERT2^ (Tg(*Vil*-cre/ERT2)23Syr), *Fos*^*TRAP2*^ (*Fos*^*tm2.1(icre/ERT2)Luo*^/J, Jackson #030323) *Tph1*^flox^ (*Tph1*^*tm1Kry*^, gift of G. Karsenty), *Htr3a*^Cre^ (Gift of N. Heintz), Glp1r^tm1Ddr^ or *Glp1r*^−/−^ (gift of D. Drucker and generously provided by J. Ayala), *Gpr43*^−/−^ (Gift of N. Arpaia), *Gpr43*^−/−^/*Gpr109a*^−/−^ (gift of S. Mehandru), *Gpr41*^−/−^ (gift of J. Gordon and M. Yanagisawa, generously provided by J. Pluznick). Gnotobiotic mice used: Germ-Free (GF) C57BL/6, *Snap25*^RiboTag^, SFB-mono-colonized, ASF-colonized (Rockefeller University), GF C57BL/6J (gift of J. Faith), GF C57BL/6J 6J and *Clostridium spp.* colonized mice (gift of D. Artis and G. Sonnenberg). Mice were bred within our facility to obtain strains described and were 7-12 weeks of age for all experiments unless otherwise indicated. For comparisons to GF mice, mice were maintained on sterilized Autoclavable Mouse Breeder Diet (5021, LabDiet, USA), the same used in the gnotobiotic facility. Female mice were used for all sequencing experiments. Male and female mice were used for all other experiments. Animal care and experimentation were consistent with NIH guidelines and approved by the Institutional Animal Care and Use Committee (IACUC) at The Rockefeller University.

### Tamoxifen treatment

For Villin^*Tph1*^ mice 1mg of tamoxifen (Sigma) was injected i.p. on two consecutive days. Mice were then analysed 2 weeks following the last dose of tamoxifen. For *Advillin*^CreERT2^ strains, mice were given 1mg of tamoxifen by i.p. injection on five consecutive days. Mice were then analyzed at least 1 week following the final dose of tamoxifen.

### Tph1-flox recombination PCR

DNA was extracted from the epithelial fraction of cells made by Percoll gradient of homogenized colon from Villin^Δ*Tph1*^ mice two weeks following tamoxifen administration using Quick Extract (Lucigen) DNA extraction buffer. Target sequences were amplified using the following primers: Tph1-forward 5’-GGATCCTAACCGAGTGTTCC-3’ Thp1-reverse-flox: 5’-GCACACCACCAACTCTTTCC-3’ Tph1-reverse-recombined: 5’-CTTGGAAGGTTTTGTATCACC-3’ PCR products were run on a 2% agarose gel and bands were analysed for the presence of the recombined band^50^.

### Controlling for stress

Due to the sensitivity of the sympathetic nervous system to stress, the following steps were taken eliminate this potential confounding factor. Experiments were not performed on days when cage changing took place. Mice were transported to the lab and sacrificed immediately, and all experiments with injections were done after a minimum of 5 days of i.p./handling habituation.

### SFB colonization

Mice mono-colonized with segmented filamentous bacteria (SFB) were kept in GF isolators and originally colonized by gavage with faecal extract from SFB mono-colonized mice kept at NYU (Littman lab). SFB colonization was verified by real time PCR using SFB-specific 16S primers; GF feces served as a negative control, Taconic Farms C57BL/6 faeces as a positive control.

### *Clostridium* spp. Colonization

Mice were colonized with *Clostridium spp.*^12^ in the Cornell Weill gnotobiotic facility.

### Altered Schaedler Flora colonization

C57BL/6 mice were maintained in germ-free conditions in ISOcage biocontainment isolator cages (Tecniplast, PA, USA) in the gnotobiotic facility at Rockefeller University. ASF colonization was achieved by inoculating germ-free mice with cecal contents of ASF donor mice stably colonized by vertical transmission (kindly provided by Amanda Ramer-Tait, University of Nebraska-Lincoln). Ceca were prepared by homogenization through a 100 µm filter in sterile phosphate-buffered saline (PBS) at a ratio of one cecum per 1 ml of PBS. Mice received 200 µl of ASF inoculum via oral-gavage twice, one week apart. The presence of members of the ASF microbial community was confirmed by a real-time PCR-based assay previously described^51^. All mice were analysed at least four weeks post colonization, with colonization further confirmed by 16s RNA sequencing of both faeces and ceca of analysed mice. We could detect RNA of at least 6 species by qPCR and 16s RNA sequencing after colonization.

### Antibiotic treatments

Broad spectrum antibiotics (0.25 g Vancomycin, 0.25 g metronidazole, 0.5 g ampicillin, and 0.5 g neomycin) were dissolved in 500 mL of filtered water and supplemented with 5 g Splenda. Individual antibiotics (0.25 g vancomycin, 0.25 g metronidazole, 0.5 g ampicillin and 0.5 g neomycin) were dissolved in 500 mL of filtered water and supplemented with 5 g Splenda. To control for the sweet taste of the antibiotic solution, 5 g of Splenda was dissolved in filtered water. Water controls were given filtered water as their drinking water. All solutions were passed through a SteriCup 0.22 µm filter. Streptomycin was prepared in sterile DPBS at a concentration of 200 mg/mL and then filtered with a 0.22 µm (EMD Millipore PES Express) syringe filter. A dose of 20 mg was given as an oral gavage of 100 µL of this stock solution.

### Tributyrin treatment

Tributyrin (Sigma W222305) was filter sterilized through a 0.22 µm (EMD Millipore PES Express) syringe filter prior to oral gavage or i.p. injection. 200 µL of tributyrin was given by oral gavage^8^. Two oral gavages were given over a period of 24 h, with the second dose given 8 h before sacrifice.

### Exendin treatment

Exendin-4 (Sigma E7144) was dissolved in sterile 0.9% saline and aliquots were kept at -20°C. 20 µg/kg Exendin-4 or saline was given by i.p. injection. Tissue was isolated 4 hours post-injection or motility was measured 2 minutes following injection.

### PYY treatment

PYY (Sigma P1306) was dissolved in sterile 0.9% saline and aliquots were kept at - 20°C. 50 µg/kg PYY or saline was given by i.p. injection. Tissue was isolated 4 hours post-injection.

### DREADD agonist

Water soluble Compound 21 (HelloBio HB6124) was dissolved in sterile 0.9% saline and aliquots were stored at -80°C. Mice were given intraperitoneal injection at a dose of 10 mg/kg or 1mg/kg.

### Tracing injections

Mice were anesthetized with 2% isoflurane with 1% oxygen followed by 1% isoflurane with 1% oxygen to maintain anaesthesia. After shaving and sterilization of the abdomen, mice were placed on a sterile surgical pad on top of a heating pad and covered with a sterile surgical drape. Ophthalmic ointment was placed over the eyes to prevent dehydration and the incision site was sterilized. Upon loss of recoil paw compression, a midline incision was made through the abdominal wall exposing the peritoneal cavity. For injections into the CG-SMG or duodenum an additional incision was made laterally to allow for better access. The duodenum, ileum, colon, or CG-SMG were located and exposed for injection. All injections were made with a pulled glass pipette using a Nanoject III. Following injection, the abdominal wall was closed using absorbable sutures and the skin was closed using surgical staples. Antibiotic ointment was applied to the closed surgical site and mice were given 0.05 mg/kg buprenorphine every 12 h for 2 days.

### Stereotactic Surgery

Mice were anesthetized using isoflurane, with induction at 4% and maintenance at 1.5-2%. Coordinates were identified using the Paxinos mouse brain atlas. For tracing studies of the nucleus tractus solitarius (NTS), mice were bilaterally injected with 150 nL of rgAAV-FLEX-CAG-tdTomato (Addgene #28306-AAVrg) into the NTS at AP -7.2, DV -4.3, ML 0.35 relative to bregma. For tracing studies of the LPGi/RVLM, mice were bilaterally injected with PRV-GFP at -6.35 AP, ML 0.9, DV -6.0. For chemogenetic activation studies, VGAT^Cre^ mice were injected with 50nL of AAV5-hSyn-DIO-hM3D(Gq)-mCherry virus into the gigantocellular reticular nucleus (Gi) at AP-6.35, DV -5.8, ML 0.5 relative to bregma. VGLUT2^Cre^ mice were injected with AAV5-hSyn-DIO-hM3D(Gq)-mCherry into the rostral ventrolateral medulla (LPGi/RVLM) at -6.35 AP, ML 0.9, DV -6.0 relative to bregma. Skin was closed using sutures.

### Subdiaphragmatic vagotomy (sdVx)

Mice were anesthetized using isoflurane (induction: 2% isoflurane with 1% oxygen, maintenance: 1% isoflurane with 1% oxygen). After shaving and sterilization of the abdomen, mice were placed on a sterile surgical pad on top of a heating pad and covered with a sterile surgical drape. Ophthalmic ointment was placed over the eyes to prevent dehydration and the incision site was sterilized. Adequate depth of anaesthesia was confirmed by loss of recoil paw compression. A midline abdominal incision was then made along the *linea alba*, exposing the peritoneal cavity. The liver was retracted using sterile, saline-dampened cotton Q-tips. The right and left vagus nerve were visualized along the oesophagus below the diaphragm by a surgical microscope and cut using microscissors. This included the hepatic, gastric, ventral and dorsal vagal trunks. 0.5 µL of CTB555/594 was then injected into the stomach or ileum to confirm successful vagotomy by a lack of labelling in the nodose ganglion. Following the procedure, the abdominal wall was closed using absorbable sutures and the skin was closed using surgical staples. Antibiotic ointment was applied to the closed surgical site and mice were given 0.05 mg/kg buprenorphine every 12 h for 2 days. For sham-operated animals the vagus nerve was similarly exposed but not cut.

### PRV tracing with sdVx

Mice were given sdVx as described above, followed by injection of a 1 µL mixture of (0.5 µL PRV-152 and 0.5 µL CTB594) into the proximal colon. The inclusion of CTB was necessary to confirm successful vagotomy and to ensure that nodose neurons observed came from retrograde transport of PRV originating in the CG-SMG and traveling through the CNS. Antibiotic ointment was applied to the closed surgical site and mice were given 0.05 mg/kg buprenorphine every 12 h for 2 days. For sham-operated animals the vagus nerve was similarly exposed but not cut.

### Whole mount intestine immunofluorescence

Briefly, mice were sacrificed by cervical dislocation and the small intestine was removed and placed in HBSS Mg^2+^Ca^2+^(Gibco) + 5% FCS. The intestine was cut open longitudinally and the luminal contents washed away in DPBS. The muscularis was then carefully dissected away from the underlying mucosa in one intact sheet. The tissue was pinned down in a plate coated with Sylgard and then fixed O/N with 4% PFA with gentle agitation. After washing in DPBS whole mount samples were then permeabilized first in 0.5% Triton X-100/0.05% Tween-20/4 μg heparin (PTxwH) for 2 hours at room temperature (RT) with gentle shaking. Samples were then blocked for 2 h in blocking buffer (PTxwH with 5% bovine serum albumin/5% donkey/goat serum) for 2 hr at RT with gentle agitation. Antibodies were added to the blocking buffer at appropriate concentrations and incubated for 2 days at 4°C. After primary incubation the tissue was washed 4 times in PTxwH and then incubated in blocking buffer with secondary antibody at concentrations within the primary antibody range for 2 hours at RT. Samples were again washed 4 times in PTxwH and then mounted with FluoroMount G on slides with 1 ½ coverslips. Slides were kept in the dark at 4°C until they were imaged.

### Restraint stress

For restraint stress experiments, each mouse was placed in a properly ventilated 50 mL conical plastic tube for 15 min. The mice could rotate but could not turn head to tail.

### Cholera toxin tracing

Mice were anesthetized and operated on as described above. 1.5 uL of 1% CTB 488, 555, or 647 in PBS with 0.1% FastGreen was injected with a pulled glass pipette using a Nanoject III into the ileum, duodenum, colon and celiac-superior mesenteric ganglion. For triple labelling, 0.5uL of 1% CTB488, 555, or 647 was injected into the duodenum, ileum, and proximal colon of the same mice. The tissue was carefully washed several times with PBS to prevent possible spill over of tracer to other tissues. Relevant tissues were then dissected after a minimum of 2-4 days post-injection.

### Ileal denervation to confirm cholera toxin tracing specificity

Mice were anesthetized as described above and the ileal vein and artery to the distal ileum was identified. A cauterizer was used to sever the main ileal artery/vein and surrounding nerves. Once the mesentery was resected, 0.5uL of CTB594 with 0.1% FastGreen was injected with a pulled glass pipette using a Nanoject III was injected into the distal ileum between the two most distal lymph nodes. The tissue was carefully washed several times with PBS to prevent possible spill over of tracer to other tissues. LNG, RNG, and CG-SMG tissues were then dissected after 2 days post-injection followed by overnight fixation in 4% PFA. Sham was identical to the previous procedure, but the mesentery was left intact.

### Nodose ganglion injection

Mice were anesthetized as described above and the ventral neck surface was cut open. Associated muscle was removed by blunt dissection to expose the trachea. The NG was located by following the vagus nerve along the carotid artery to the base of the skull. Fine forceps were used to separate the vagus from the carotid artery, and the NG body was exposed by careful dissection. 1uL of AAV9-Syn-ChrimsonR-tdTomato (Addgene #59171-AAV9) with 0.1% FastGreen was injected with a pulled glass pipette using a Nanoject III. The skin was then closed with absorbable sutures and antibiotic cream was applied. Relevant tissues were dissected at a minimum of 2 weeks post-injection.

### FosTRAP2-PRV analysis

*Fos*^TRAP2^ mice were crossed with *Rosa26*^lsl-tdTomato^:RiboTag mice to generate a TRAP2^Tom:RiboTag^ reporter strain. Mice were habituated and singly housed for at least 5 days. A single oral gavage of streptomycin (20mg/mouse) was given and after 24 hours mice were injected i.p. with an aqueous solution 4 hydroxytamoxifen (Deisseroth paper) to induce Cre recombinase activity in *Fos* expressing cells, resulting in tdTomato and RPL22-HA expression by recently activated neurons. After 7 days, we injected 1uL of PRV-152 (GFP) into the proximal colon. Four days later, were perfused and relevant brain areas were analysed for the presence of Tomato+ and GFP+ neurons.

### CTB NG and CG-SMG counting

CTB488 was injected into the duodenum, ileum, and proximal colon. Mice were sacrificed by cervical dislocation and the CG-SMG and NG were harvested and fixed overnight in 4% PFA. Tissue was then washed four times in DPBS at RT and permeabilized in PTxwH for 4 h at RT. Primary antibody anti-AlexaFluor488 (1:400, Thermo Fisher Scientific, A-11094) was added to the samples in PTxwH and incubated at 4°C for 48 h. Samples were washed four times in PTxwH at RT and then stained with goat-anti rabbit AF555/568/647 at 4°C for 24 h. Samples were washed four times in PTxwH at RT, covered in Fluormount G, and coverslipped for confocal imaging. Each ganglion was captured in full by multiple z-stacks and the total number of CTB+ neurons were counted.

### Celiac ganglion tracing

Mice were anesthetized and operated on as described above. 1.5 uL of AAVrg-hSyn1-Cre with 0.1% FastGreen was injected into the CG-SMG of *Rosa26*^lsl-tdTomato^ mice. 1.5 uL of AAV2-CAG-FLEX-tdTomato with 0.1% FastGreen was injected into the CG-SMG of *Snap25*^cre^ mice. Intestine samples were dissected after 2.5 weeks for AdipoClear, RIMS, or Focus Clear analysis.

### Viscerofugal tracing

Mice were anesthetized and operated on as described above. 1.5 uL of AAV6-CAG-FLEX-tdTomato with 0.1% FastGreen was injected into the ileum wall of *Snap25*^cre^ mice. CG-SMG samples were then dissected after 2.5 weeks for whole mount immunofluorescence analysis.

### Virus

All viruses used in these studies were: AAV5-hSyn-DIO-mCherry (Addgene), AAVrg-CAG-tdTomato (Addgene), AAVrg-CAG-FLEX-tdTomato (Addgene), AAV6-CAG-FLEX-tdTomato (Addgene), AAVrg-hSyn1-Cre (Janelia), AAV2-hSyn-hM3Dq-mCherry (Addgene), AAV2-hSyn-hM4Di-mCherry (Addgene), AAV9-Syn-ChrimsonR-tdTomato (Addgene #59171-AAV9), PRV-152 (Gift of L. Enquist), PRV-614 (Gift of L. Enquist). Fast Green (Sigma) was added (0.1%) to virus injected into peripheral tissues.

### Fluorogold Labelling

A stock solution of 4 mg/mL Fluorogold (Fluorochrome) was made in sterile 0.9% saline and then filter sterilized through a 0.22 µm syringe filter. An i.p. injection of 300 µL of Fluorogold solution was given 3 days before tissue harvesting.

### Retrograde PRV Tracing

Mice were anesthetized and operated as described above. PRV Bartha 152 (GFP) or 614 (RFP) were a gift of L. Enquist. 3uL with 0.1% FastGreen was injected with a pulled glass pipette using a Nanoject III into the wall of the ileum, duodenum, and colon. Brains and spinal columns were harvested three and four days after injection.

### Chemogenetics of CG-SMG neurons

1 µL of AAV2-hSyn-hM3Dq-mCherry or AAV2-hSyn-hM4Di-mCherry (Addgene) was injected into the CG-SMG of C57BL/6J mice. Mice were then sutured, and staples were applied. Antibiotic ointment was applied to the closed surgical site and mice were given 0.05 mg/kg buprenorphine every 12 h for 2 days. After 2 weeks mice were habituated to i.p. injections for 5 days before administration of 1 mg/kg or 10mg/kg Compound 21.

### Antibodies

The following primary antibodies were used, and unless otherwise indicated concentrations apply to all staining techniques: NeuroTrace AF569/AF647 (1:200, Thermo Fisher Scientific, N21482/N21483), GFP Tag AF488/555/594/647 (1:400, Thermo Fisher Scientific A21311/A31851/A21312/A31852), TH (1:200, Aves Labs, TYH; 1:400, Millipore Sigma, AB152; 1:200 Millipore Sigma, AB1542), BIII-Tubulin (1:400, Millipore Sigma, T2200; 1:200, Aves Labs, TUJ), NPY (1:200, Immunostar, 22940), RFP (1:200, Sicgen, AB8181; 1:200 and 1:1000 AdipoClear brain, Rockland, 600-401-379), ANNA-1 (1:200,000, Gift of Dr. Vanda A. Lennon), cFos (1:500, Cell Signaling Technologies, 2250S), HA (1:400, Cell Signaling Technologies, 3724S), 5-HT (1:200, Millipore Sigma, MAB352), anti-AlexaFluor488 (1:400, Thermo Fisher Scientific, A-11094), and VGLUT2 (1:200, Millipore Sigma, AB2251-I). Fluorophore-conjugated secondary antibodies were either H&L or Fab (Thermo Fisher Scientific) at a consistent concentration of 1:400 in the following species and colors: goat anti-rabbit (AF488/568/647), goat anti-rat (AF488/647), goat anti-chicken (AF488/568/647), goat anti-human (AF568/647), donkey anti-guinea pig (AF488/647), donkey anti-rabbit (AF568/647), donkey anti-goat (AF568/647), donkey anti-sheep (AF568/790).

### Intestine dissections

Mice were sacrificed and duodenum (1 cm moving proximal from the gastroduodenal junction), ileum (1 cm moving proximal from the ileocecal junction), or colon (4 cm moving proximal from the rectum) was removed. For AdipoClear fecal contents were flushed from the lumen and tissue was left intact. Tissue used for RIMS or FocusClear were cut open longitudinally and fecal contents were washed out. For dissection of the muscularis, following the above procedures, the intestinal tissue was placed on a chilled aluminium block with the serosa facing up^14^. Curved forceps were then used to carefully remove the muscularis.

### Nodose ganglion (NG) dissections

Mice were sacrificed and the ventral neck surface was cut open. Associated muscle was removed by blunt dissection to expose the trachea. The NG was then located by following the vagus nerve along the carotid artery to the base of the skull. Fine scissors were used to cut the vagus nerve below the NG and superior to the jugular ganglion.

### Celiac-superior mesenteric ganglion dissections

Mice were sacrificed and a midline incision was made, and the viscera were reflected out of the peritoneal cavity. The intersection of the descending aorta and left renal artery was identified, from which the superior mesenteric artery was located. The CG-SMG is wrapped around the superior mesenteric artery and associated lymphatic vessels. Fine forceps and scissors were used to remove the CG-SMG.

### Dorsal root ganglion dissections

The spinal column was isolated, cleaned of muscle, and bisected sagittally. The spinal cord was removed leaving the dorsal root ganglion (DRG) held in place by the meninges. The thoracic 13 DRG was identified by its position just caudal to thoracic vertebra. The meninges were cleared and individual DRGs were removed with fine forceps and scissors.

### Spinal cord dissections

For 3DISCO analysis the spinal cord was isolated by hydraulic extrusion as previously described^52^. For whole spinal column calAdipoClear imaging, the entire spinal column was removed with associated tissue. Costae and muscle were carefully trimmed to reduce the size of the sample to fit into a 5 mL Eppendorf tube while avoiding disrupt ventral tissue attached to the spinal cord.

### RiboTag

Heterozygous or homozygous *Snap25*^RPL22HA^ were used for TRAP-seq analysis as no differences were found between either genotype. For NG, DRG, and CG-SMG IP, tissues were isolated as described above. The RiboTag IP protocol was then followed (http://depts.washington.edu/mcklab/RiboTagIPprotocol2014.pdf) with the following modifications. All samples were homogenized by hand with a dounce homogenizer in 2.5 mL supplemented homogenization buffer (changes per 2.5 mL: 50 µL Protease Inhibitor, 75 µL heparin (100 mg/mL stock), 25 uL SUPERase· In RNase Inhibitor). Samples were then centrifuged for 10 minutes at 10,000 G, after which 800 µL of supernatant was removed and 5 µL of anti-HA antibody (Abcam, ab9110) was added. Samples were kept rotating at 4°C with antibody for 1 hour. 200 µL of Thermo Protein magnetic A/G beads were washed with homogenization buffer, added to the sample, and kept rotating for 30 minutes at 4°C. The beads were washed four times with high-salt buffer and samples were eluted with 100 µL of PicoPure lysis buffer. RNA was extracted using the Arcturus PicoPure RNA isolation kit (Applied Biosystems) according to the manufacturer’s instructions.

### RiboTag RNA-sequencing

RNA libraries were prepared using SMARTer Ultra Low Input RNA (ClonTech Labs) and sequenced using 75 base-pair single end reads on a NextSeq 500 instrument (Illumina). Reads were aligned using Kallisto^53^ to the Mouse transcriptome (Ensembl, release v91). Transcript abundance files were then used in the DESeq2 R package^54^, which was used for all downstream differential expression analysis and generation of volcano plots. For intestine samples cre+ samples were compared with cre-samples to generate a list of immunoprecipitated (IP) enriched genes (log2FC > 1 and padj < 0.05). This IP enriched list was then used to perform downstream analysis. Differentially expressed genes between samples were defined as those contained within the total IP enriched list from tissues being compared and with a cutoff of log2FC > 1. PCA plots were generated from log transformed DEseq2 data, as indicated in figure legends, with the FactoMineR R package. GSEA pre-ranked analysis was performed with desktop software and the C5 gene ontology database using 1000 permutations. Gene ontology enrichment analysis was performed with differentially expressed genes (log2FC > 1, padj < 0.05) using the TopGO R package and a Fisher test with an elimination algorithm was used to calculate significance.

### 16S sample processing

16s samples were processed utilizing a Promega Maxwell® RSC 48 Instrument. Following DNA extraction from all samples, DNA samples were quantified using a ThermoFisher Quant-It dsDNA High-Sensitivity Kit on a microplate reader.

### 16S sequencing

16s sequencing was performed on either the Illumina HiSeq 2500 or the Illumina MiSeq depending on project-specific needs. Raw paired-end fastq files containing sequence reads were merged at the overlapping region to produce a single 16s contig. All merged sequences having more than 1 expected error per read were filtered. Operational taxonomic units (OTUs) were generated by clustering sequences with a 99% correspondence and chimera sequences were removed using usearch^55^ (v11). Reads were mapped against the OTU reference to generate a matrix of counts. Subsequently, OTU taxonomy and classification were performed with mothur^56^ (v1.40.5) using the greengenes database. Next, statistical analysis were implemented using the phyloseq^57^ package for R.^56^ (v1.40.5) using the greengenes database. Next, statistical analysis were implemented using the phyloseq^57^ package for R.

### Brain immunofluorescence

Mice were sacrificed and transcardially perfused with cold PBS with heparin followed by cold 4% PFA (Electron Microscopy Sciences). The intact brain was separated carefully from the skull and placed in 4% PFA, and then rotated for 48 h at 4°C. Whole brains were washed with PBS/0.03%Azide and sectioned at 50 μm on a Leica vibratome for immunofluorescence. Samples were then permeabilized in 0.5% Triton/0.05 Tween-20 in PBS (PTx) followed by blocking in 5% goat serum in PTx each for 2 h at room temperature. Primary antibody was added to the blocking buffer and samples were incubated with constant rotation at 4°C overnight. Four 15-minute washes were done in PTx at RT after which samples were moved to blocking buffer with secondary antibody. Slices were incubated in secondary antibody for 2 hours at room temperature followed by four 15-minute washes in PTx at room temperature. Samples were then placed on microscope slides, covered in Fluormount G, and coverslipped.

### Confocal imaging

Whole mount intestine, NG, DRG, and CG-SMG samples were imaged on an inverted LSM 880 NLO laser scanning confocal and multiphoton microscope (Zeiss) and on an inverted TCS SP8 laser scanning confocal microscope (Leica).

### PRV counting

Images of brainstem vibratome slices were taken at 10x magnification. The raphe pallidus, gigantocellular nucleus, and lateral paragigantocellular nucleus/rostral ventrolateral medulla were identified based on the Allen Brain Atlas. All VGAT^GFP^ and VGLUT2^GFP^ cells were counted within this region, as well as all PRV-RFP+ cells. These numbers were then averaged across each brainstem region for all slices from a single animal. Thus, each point on the graph is representative of one animal.

### RIMS clearing

Briefly, following secondary staining CG-SMG, nodose and DRG were submerged in Refractive Index Matching Solution (RIMS) for 30-120 min then mounted in RIMS solution on a glass slide and sealed with a coverslip for confocal imaging^29^.

### FocusClear

Whole intestine and celiac ganglion samples were first fixed in 4% PFA overnight at 4°C. Samples were then washed three times in DPBS at RT. Samples were placed into 250 μL of FocusClear solution for 15-20 minutes. They are then transferred to MountClear solution on a glass slide and a 1 ½ coverslip was used to seal the sample in place.

### 3DISCO

3DISCO clearing of whole spinal cord was done as previously described^58^.

### AdipoClear

Adipoclear whole tissue clearing was adapted from Adipoclear protocol^23^. Mice were sacrificed and intestinal sections were removed followed by overnight fixation in 4% PFA. Tissues were washed in PBS then dehydrated in 20/40/60/80/100% Methanol in B1N followed by dichloromethane. Tissues were then rehydrated in 100/80/60/40/20% methanol in B1N. Subsequently, samples were washed in PTxwH and then incubated in primary antibody dilutions in PTxwH for 7 Days. Samples were washed in PTxwH then incubated in secondary antibody at 1:400 in PTxwH for 7 days. Samples were again washed in PTxwH followed by PBS then dehydrated in 20/40/60/80/100% methanol followed by dichloromethane and finally cleared in dibenzyl ether.

### calAdipoClear

CalAdipoclear whole tissue clearing was adapted from Adipoclear protocol^23^. Briefly, mice were sacrificed and perfused with PBS followed by 4% PFA. Whole spinal columns were removed and put into 4% PFA overnight. Tissues were washed in PBS then dehydrated in 20/40/60/80/100% methanol in B1N followed by dichloromethane. Tissues were then rehydrated in 100/80/60/40/20% methanol in B1N. Subsequently, samples were decalcified in Morse Solution (1-part 45% formic acid/1-part 0.68 mM sodium citrate dihydrate) overnight followed by PTxwH washes. Samples were then incubated in primary antibody dilutions in PTxwH for 7 Days. Samples were washed in PTxwH then incubated in secondary antibody at 1:400 in PTxwH for 7 days. Samples were again washed in PTxwH followed by PBS then dehydrated in 20/40/60/80/100% methanol followed by dichloromethane and finally cleared in dibenzyl ether.

### Light sheet microscopy and 3D reconstructions

Whole-tissue cleared samples were imaged submerged in DBE on a LaVision Biotech Ultramicroscope II with 488 nm, 561nm, 640 nm, or 785 light sheet illumination using a 1.3x or 4x objective with 2.5um Z-slices. Images were adjusted post hoc using Imaris x64 software (version 9.1 Bitplane) and 3D reconstructions were recorded as mp4 video files. Optical slices were taken using the orthoslicer or oblique slicer tools.

### ClearMap analysis

All analyses for whole-brain studies were performed adapting ClearMap pipeline (latest version available from www.idisco.info, see also^30^).

### Quantification of iEAN

A minimum of 10 images were randomly acquired across a piece of whole mount muscularis. These images were then opened in ImageJ, and the cell counter was used to count the number of ANNA-1+ cells in a given field. This number was then multiplied by a factor of 2.95 (20x objective) or 3.125 (25x objective), to calculate the number of counted neurons per square millimeter (mm^2^). The average of 10 (or more) images were then calculated and plotted. Thus, every point on a given graph corresponds to a single animal.

### Quantification of CG-SMG cFos

Mice were sacrificed by cervical dislocation and CG-SMG were harvested and fixed overnight in 4% PFA. CG-SMG were then washed four times in DPBS at RT and permeabilized in PTxwH overnight RT. Primary antibody cFos (1:500, Cell Signaling Technologies, 2250S) was added to the samples in PTxwH and incubated at RT for 48 h. Samples were washed four times in PTxwH at RT and then stained with goat-anti rabbit AF555/568/647 at RT for 48 h. Samples were washed four times in PTxwH at RT, covered in Fluormount G, and coverslipped for confocal imaging. We first established criteria for identifying neuronal cFos+ nuclei by staining CG-SMG from restraint-stressed mice, a condition known to activate the sympathetic nervous system^10,26^. FluoroGold was used to identify sympathetic neurons and cFos+ nuclei were defined as morphologically circular with a diameter of 8-14um. These criteria were sufficient to distinguish between small intensely fluorescent cells and possibly macrophages that also have cFos expression. We captured all sympathetic neurons within the CG-SMG, as defined by tyrosine hydroxylase staining, FluoroGold fluorescence, tdTomato fluorescence, or autofluorescence (experiment dependent), with multiple z-stack images. All images were analyzed in Image-J. Total cFos+ nuclei were counted using the Cell Counter plugin for Image-J, and data were not normalized to area or volume. Each data point represents the number of cFos+ cells per CG-SMG.

### Brainstem cFos counting experiments

All mice (GF, exGF, SPF, SNS^hM4Di^, and VGLUT2^RiboTag:tdTomato^) were fasted for 18-24 hours before perfusion (exGF, SPF, and VGLUT2^RiboTag:tdTomato^) or injection of Compound 21 (SNS^hM4Di^). Mice were then perfused 3 hours post Compound 21 injection or 24 hours post streptomycin gavage. Brains were sectioned as described above and sections were permeabilized in 0.5% Triton/0.05 Tween-20 in PBS (PTxwH) followed by blocking in 5% goat serum in PTxwH each for 2 h at room temperature. cFos primary antibody was added to the blocking buffer and samples were incubated with constant rotation for 48 hours at 4°C. Four 15-minute washes were done in PTxwH at RT after which samples were moved to blocking buffer with secondary antibody. Slices were incubated in secondary antibody for 2 hours at room temperature followed by four 15-minute washes in PTxwH at room temperature. Samples were placed on microscope slides, covered in Fluormount G, and coverslipped. All sections containing NTS/AP were imaged and included in counting of cFos+ cells. Therefore, each data point represents the total number of cFos+ cells per relevant brain area captured.

### RNAScope

Nodose ganglia (NG) from C57Bl/6 mice were dissected as described above. Once removed ganglia were dipped in Fast Green (1%, Sigma-Aldrich) to assist with visualization when slicing and flash frozen in OCT. 15 um sections of NG were sliced on a cryostat for RNAScope. Samples were processed and stained with Scn5a, positive control or negative control probes according to the manufacturer’s instructions. Samples were mounted in Prolong gold antifade with DAPI (Thermo-Fisher) for imaging and imaged within 24 hours on an inverted LSM 880 NLO laser scanning confocal and multiphoton microscope (Zeiss) and images were processed using Image J.

### RNAScope/IHC

C57Bl/6 mice were injected bilaterally with CTB 488 into the colon as described above. NG were dissected 1 week post injection as described above, dipped in Fast Green (1%, Sigma-Aldrich) to assist with visualization and flash frozen in OCT. 15 um sections of NG were sliced on a cryostat for RNAScope/IHC. Samples were processed and stained with Scn5a, positive or negative control probes according to the manufacturer’s instructions. After in situ hybridization sections were washed three times in wash buffer (1X, ACDBio) and then fixed in 1% PFA in TBS for 10 minutes at 4 C to stabilize the ISH labeling. Samples were next washed three times in TBS-T and incubated in 10% Goat Serum in TBS with 1% BSA for 30 minutes. Samples were stained with anti Alexafluor-488 antibody (1:500, Thermo-Fisher) for 1 hour in TBS-1% BSA. After primary antibody staining, sections were washed three times for 5 minutes each in TBST and stained with Goat anti rabbit AF488 (1:500, Thermo-Fisher) in TBS-1% BSA for 30 minutes. Samples were again washed three times for 5 minutes each in TBST and finally mounted in Prolong gold antifade with DAPI (Thermo-Fisher) for imaging. Slides were imaged within 24 hours of mounting on an inverted LSM 880 NLO laser scanning confocal and multiphoton microscope (Zeiss) and images were processed using Image J.

### Intestine motility measurements

For measurement of total intestinal transit time, mice were given an oral gavage of 6% carmine red dissolved in 0.5% methylcellulose (made with sterile 0.9% saline). Total intestinal transit time was measured as the time from oral gavage it took for mice to pass a fecal pellet that contained carmine. To measure colonic motility a glass bead (3 mm diameter) was pushed into the colon to a distance of 2 cm from the anal verge. The time required for expulsion of the glass bead was measured and taken as an estimate of colonic motility. Mice in both assays were injected 2 minutes before starting with i.p. Compound 21 (1mg/kg or 10mg/kg as indicated).

### Open Field Test

For locomotion activity assessment an open field test (OFT) was performed using a small cubic box, measuring 27.3 cm^3^. The top of the cube of the OFT box is uncovered. Mice were placed in the bottom surface, and movements were recorded over the course of 2 sessions of 5 minutes. The first session was measured without manipulating the animal (basal). The second session was measured immediately after an I.P. injection of Compound 21 (10mg/kg). Computer-tracking program EthoVision XT (Noldus) was used to analyse the movements of the animal over time. Total distance travelled and velocity was assessed.

### *Salmonella* infections

CBA/J mice were given an oral gavage of 10^9^ WT *Salmonella* typhimurium (IR715). For all *Salmonella* infections, a single aliquot of either strain of *Salmonella* was grown in 3 ml of luria broth (LB) overnight at 37°C with agitation. Bacteria were then sub-cultured (1:300) into 3 ml of LB for 3.5 hours at 37°C with agitation, and diluted to final concentration in 1 ml of LB. Bacteria were inoculated by gavage into recipient mice in a total volume of 100 µl and mock infected mice were given an oral gavage of 100 µl LB.

### Colony forming unit counting

Faecal pellets from *Salmonella*-infected mice were weighed and then disrupted in 400 μL of DPBS. Serial dilutions were made from the original suspension and then 5μL of each dilution was plated onto Salmonella-Shigella plates. The plates were then incubated overnight, and the number of black colonies were counted for the serial dilution with the clearest delineation of single units. This number was then multiplied by the dilution factor and by 80 to give the number of colony forming units (CFUs) in the original suspension. CFU numbers were then divided by the original fecal pellet weight to give the number of CFUs per mg of faeces.

### Cecal butyrate measurements

Concentrations of acetate, propionate, and butyrate were measured as previously described^59^. Briefly murine cecal content samples were collected directly into Bead-Ruptor tubes with 2.8-mm ceramic beads (OMNI International) and immediately frozen on dry ice. After thawing, samples were extracted with 80% methanol containing internal standards of deuterated SCFA (d-3 acetate, d-5 propionate, and d-7 butyrate; Cambridge Isotope Laboratories). Pellets were resuspended at a ratio of 100 mg/ml and homogenized. Homogenized samples were centrifuged at 20,000 *g* for 15 min at 4°C. The supernatant was derivatized for 60 min at 65°C with one volume of 50 mM, pH 11.0 borate buffer, and four volumes of 100 mM pentafluorobenzyl bromide (Thermo Scientific) in acetone (Fisher Scientific). The SCFA were extracted in n-hexane and then further diluted 1:10 in n-hexane. Extracted SCFA were quantified by GCMS (Agilent 7890A GC System; Agilent 5975C MS detector) operating in negative chemical ionization mode with methane as the reagent gas. MassHunter software was used for data analysis (B07.0; Agilent Technologies).

### Statistical analysis

Significance levels indicated are as follows: *P < 0.05, **P < 0.01, ***P < 0.001. All data are presented as mean ± s.d.. All statistical tests used were two-tailed. The experiments were not randomized, and no statistical methods were used to predetermine sample size. Multivariate data was analysed by one-way ANOVA and Tukey’s multiple comparisons post hoc test. Comparisons between two conditions were analysed by unpaired Student’s t-test. We used GraphPad PRISM version 8.0d and R 3.4.3 for generation of graphs and statistics.

## Supporting information

Supplemental Annotations

## Acknowledgements

We thank all Mucida Lab members past and present for assistance in experiments, fruitful discussions and critical reading of the manuscript. In particular we thank; A. Rogoz for the maintenance of gnotobiotic mice; S. Gonzalez for maintenance of SPF mice; T. Rendon and B. Lopez for genotyping; C. Nowosad for setting up the ASF mice; A. North, C. Pyrgaki, T. Tong, C. Rico, and P. Ariel of the Rockefeller University Bio-imaging Research Center for assistance with the light sheet microscopy and image analysis; the Rockefeller University Genomics Center for RNA sequencing; Rockefeller University employees for continuous assistance; T.Y. Oliveira for valuable discussions on TRAPseq approaches and help with intial analysis; S. Poliak (Kallyope) for help in setting up tracing experiments and valuable input throughout the project; K. Loh and A. Ilanges for fruitful experimental discussions; J. Friedman and V. Ruta for the generous use of lab equipment; J. Cross, A. Pickard, and A. Rao for performing the SCFA analysis; A. Lockhart, V. Jové, and V. Ruta for critical reading of the manuscript; V. Jové for critical reading of the reviewer responses; L. Vosshall, M. Nussenzweig, G. Victora and J. Lafaille and their respective lab members for fruitful discussions and suggestions; D. Littman and M. Xu for generously providing feces from SFB mono-colonized mice; K. Honda, D. Artis, and G. Sonnenberg for generously providing GF mice and *Clostridium* consortium colonized mice; N. Arpaia for *Gpr43*^-/-^ mice; S. Mehandru for *Gpr109a*^-/-^ *Gpr43*^-/-^ mice; J. Pluznick, J. Gordon, and M. Yanagisawa for *Gpr41*^-/-^ mice; I. Chiu and J. Wood for *Nav1.8*^Cre^ mice; C. Woolf and R. Kuhner for SNS^Cre^ mice; A. Ramer-Tait for ASF ceca; J. Faith for providing GF. We finally thank R.U. Muller for his inspiration. This work was supported NIH Virus Center grant no. P40 OD010996, NIH F31 Kirchstein Fellowship (P.A.M.), NCATS NIH UL1TR001866 (P.A.M., D.M.), Philip M. Levine Fellowship (P.A.M.), Kavli Graduate Fellow (P.A.M.), Kavli Postdoctoral Fellow (M.S.), Anderson Graduate Fellowship (F.M.), Leon Levy Fellowship in Neuroscience (J.d.M.), the Leona M. and Harry B. Helmsley Charitable Trust (D.M.), the Burroughs Wellcome Fund PATH Award (D.M.), Transformative R01DK116646 (D.M.).

## Author Contributions

P.A.M. initiated, designed, performed and analysed the research, helped with supervision of the research and wrote the manuscript. M.S. performed brain AdipoClear and ClearMap analysis, provided guidance on brain experiments, analysed data, and helped with manuscript preparation. Z.K. performed experiments, analysed research, and reviewed the manuscript. P.W. performed stereotaxic brain injections, provided technical advice, and reviewed the manuscript. A.I. performed stereotaxic brain injections of the NTS and AP, provided technical advice, and reviewed the manuscript. F.M. performed experiments, analysed data, and helped with manuscript preparation. T.B.R.C. gave guidance on bioinformatic approaches, wrote analysis scripts, and performed 16S sequencing analysis. I.d.A, W.H., and M.P. gave expert advice and technical support with vagal nerve experiments including nerve recordings. J.d.M. performed electrophysiology experiments and analysed the data. D.M. initiated, designed and supervised the research, and wrote the manuscript.

## Supplementary Video Legends

**Supplementary Video 1.** AdipoClear whole mount of colon stained with anti-RFP showing tdTomato+ fibres from the CG-SMG 2 weeks post AAVrg-CAG-FLEX-tdTomato injection.

**Supplementary Video 2.** AdipoClear whole mount of cecum stained with anti-RFP showing tdTomato+ fibres from the CG-SMG 2 weeks post AAVrg-CAG-FLEX-tdTomato injection.

**Supplementary Video 3.** 3DISCO whole mount of spinal cord with neurons retrograde traced from the CG-SMG with CTB555.

**Supplementary Video 4.** calAdipoClear whole mount of spinal cord stained with anti-RFP (red) and anti-TH (green), showing the CG-SMG 4 days post PRV infection in the ileum.

**Supplementary Video 5.** calAdipoClear whole mount of spinal cord stained with anti-RFP (red) and anti-TH (white) 4 days post PRV infection in the ileum.

**Supplementary Video 6.** AdipoClear whole mount of brain stained with anti-RFP (red) 3 and 4 days post PRV infection in the ileum, showing the brainstem.

**Supplementary Video 7.** AdipoClear whole mount of duodenum stained with anti-RFP (white) 2 weeks post injection of AAV9-Syn-ChrimsonR-tdTomato into the RNG.

**Supplementary Video 8.** AdipoClear whole mount of ileum stained with anti-RFP (white) 2 weeks post injection of AAV9-Syn-ChrimsonR-tdTomato into the RNG.

**Supplementary Video 9.** AdipoClear whole mount of colon stained with anti-RFP (white) 2 weeks post injection of AAV9-Syn-ChrimsonR-tdTomato into the RNG. Zoomed in on the epithelium.

**Supplementary Video 10.** AdipoClear whole mount of colon stained with anti-RFP (white) 2 weeks post injection of AAV9-Syn-ChrimsonR-tdTomato into the RNG. Zoomed in on the epithelium.

**Supplementary Video 11.** AdipoClear whole mount of colon stained with anti-RFP (white) 2 weeks post injection of AAV9-Syn-ChrimsonR-tdTomato into the RNG. Zoomed in on a single semilunar fold.

