## Supplementary figures and images for "Microbes modulate sympathetic neurons via a gut-brain circuit"

### Extended Fig1

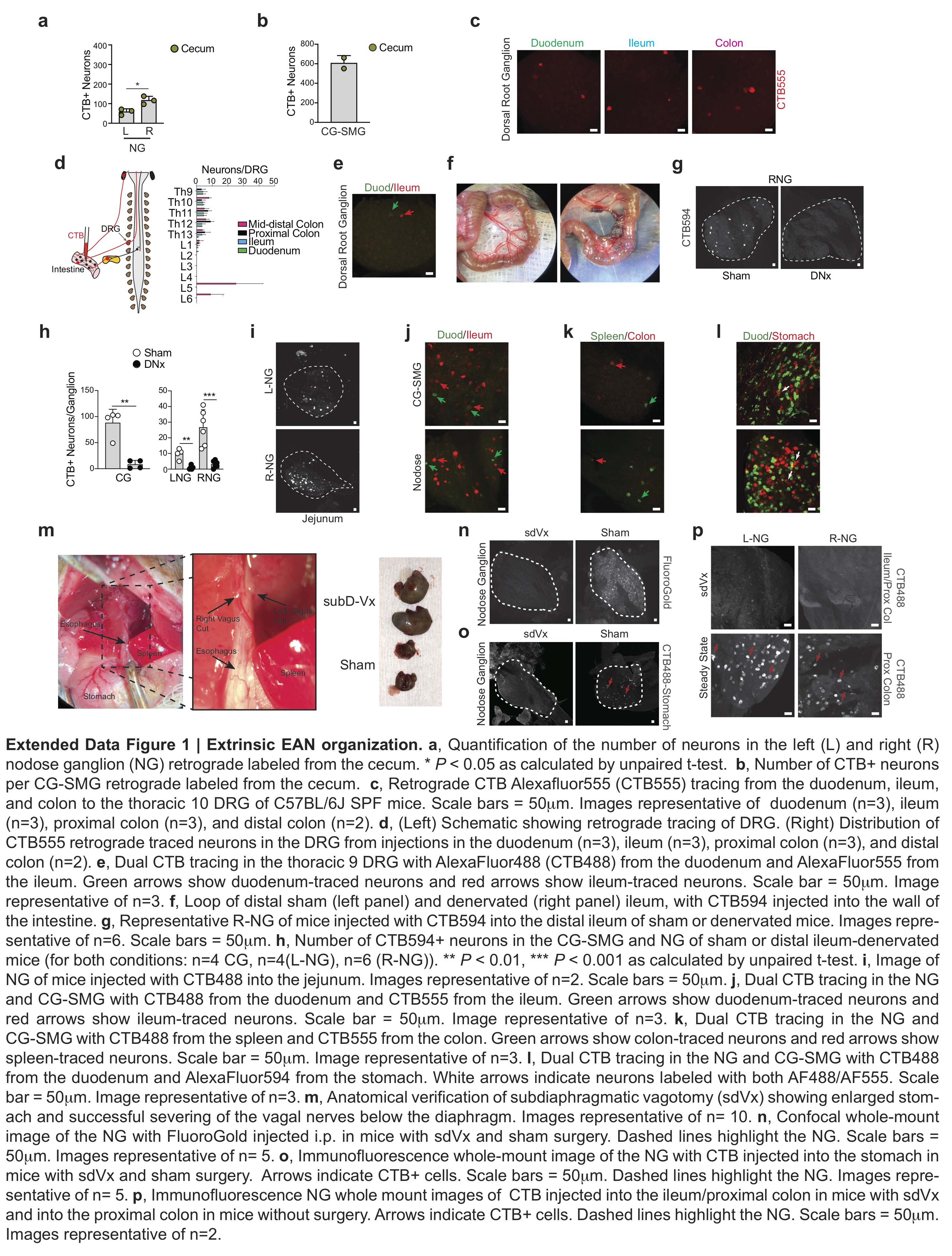

### Extended Fig2

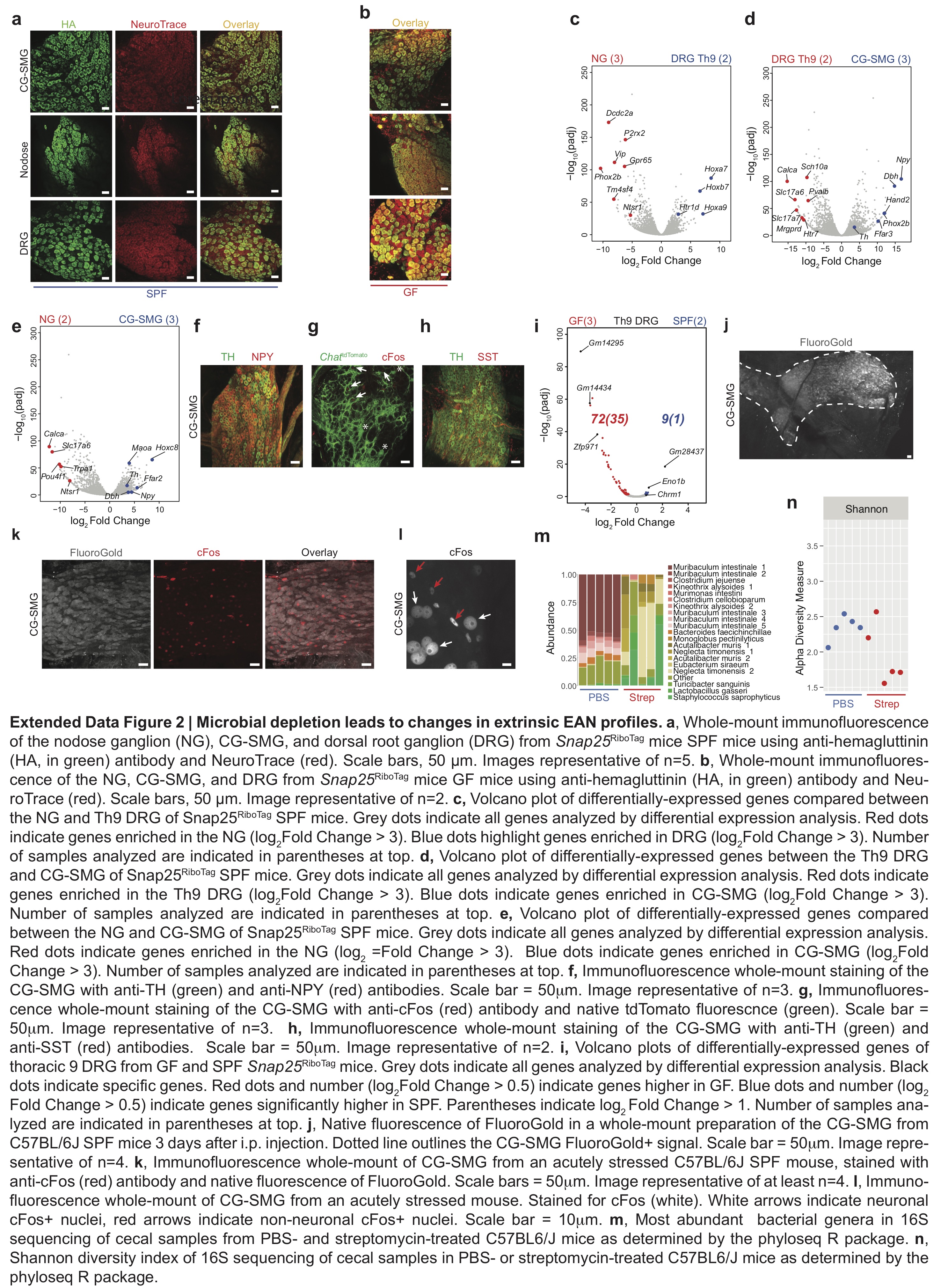

### Extended Fig3

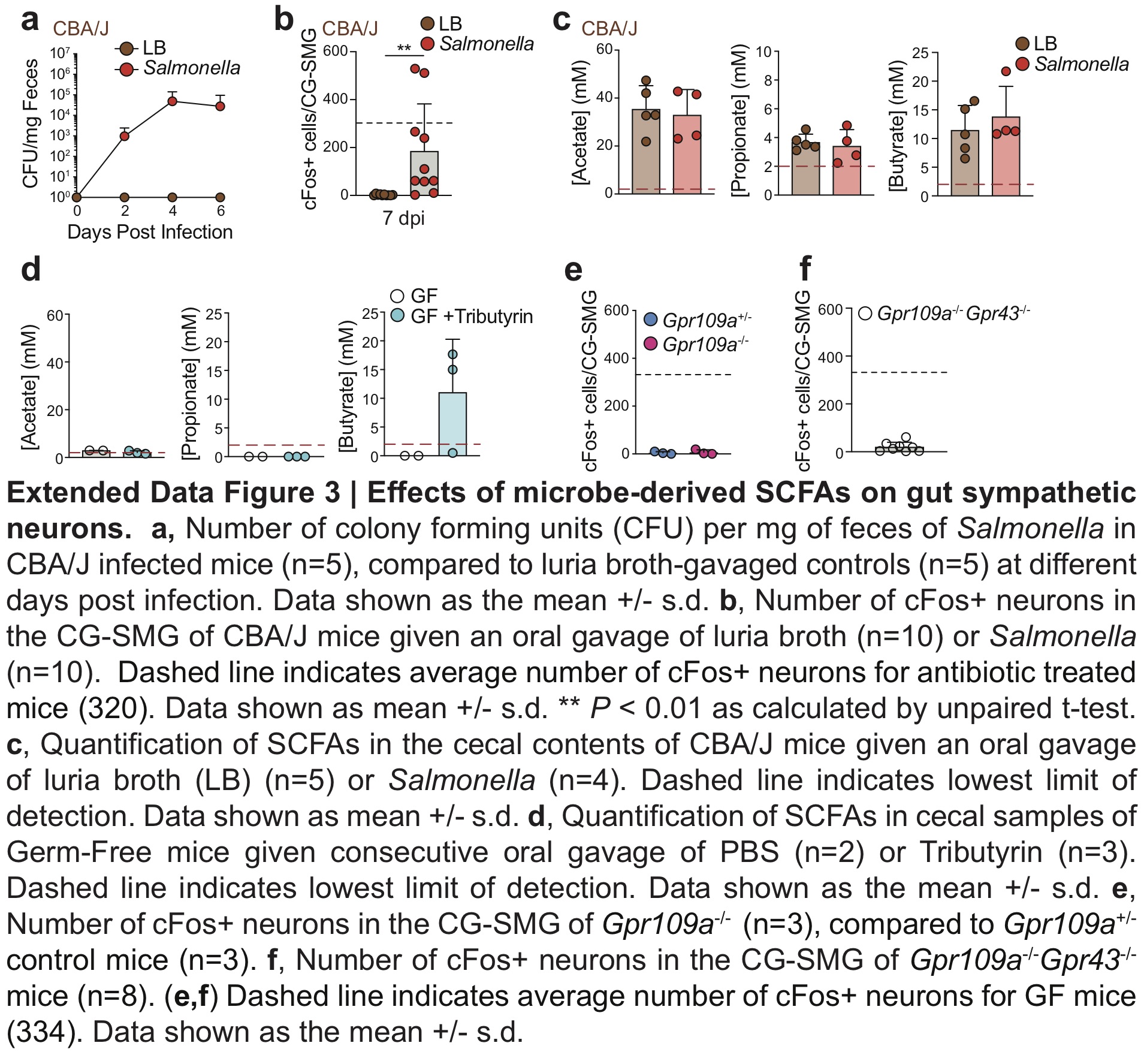

### Extended Fig4

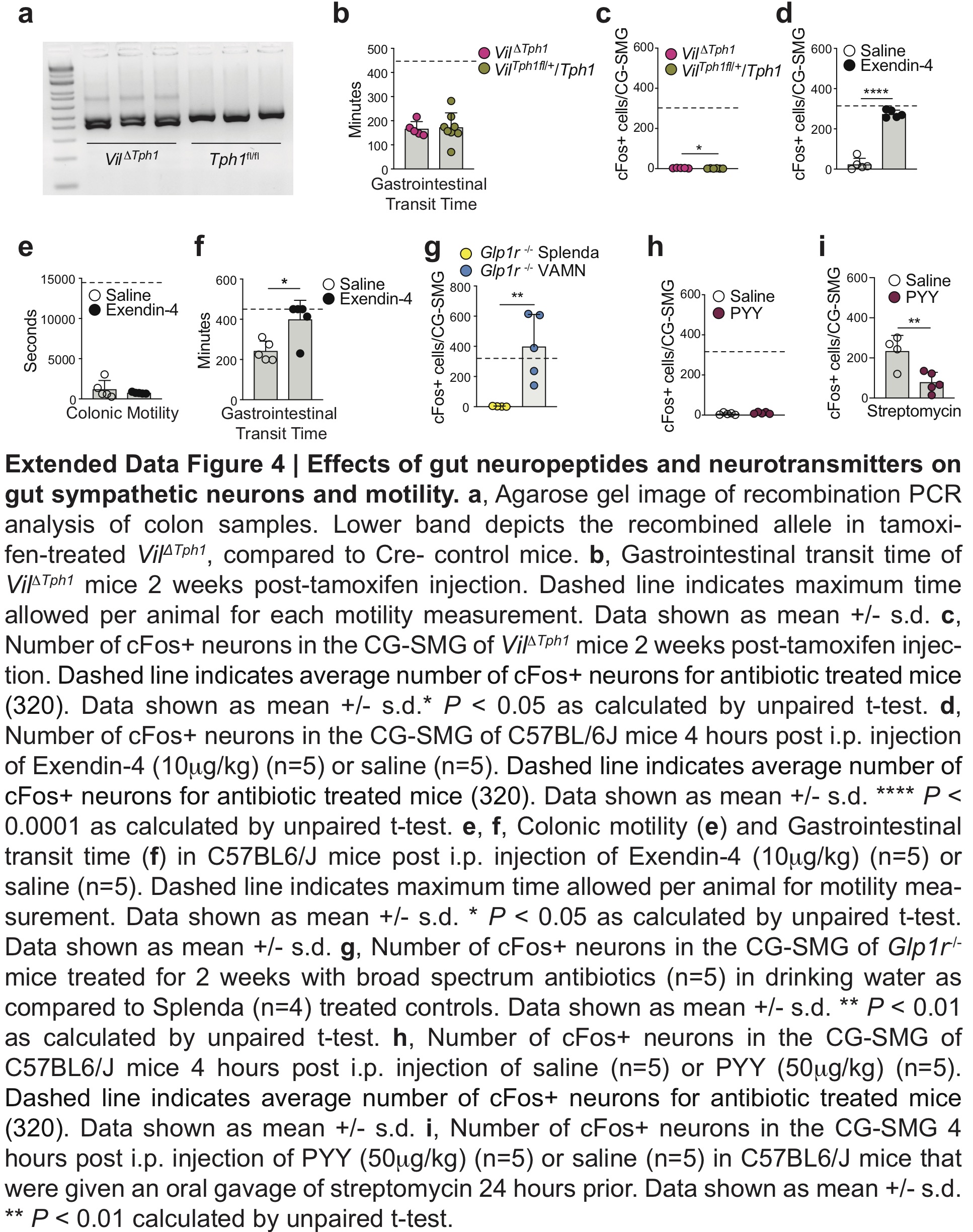

### Extended Fig5

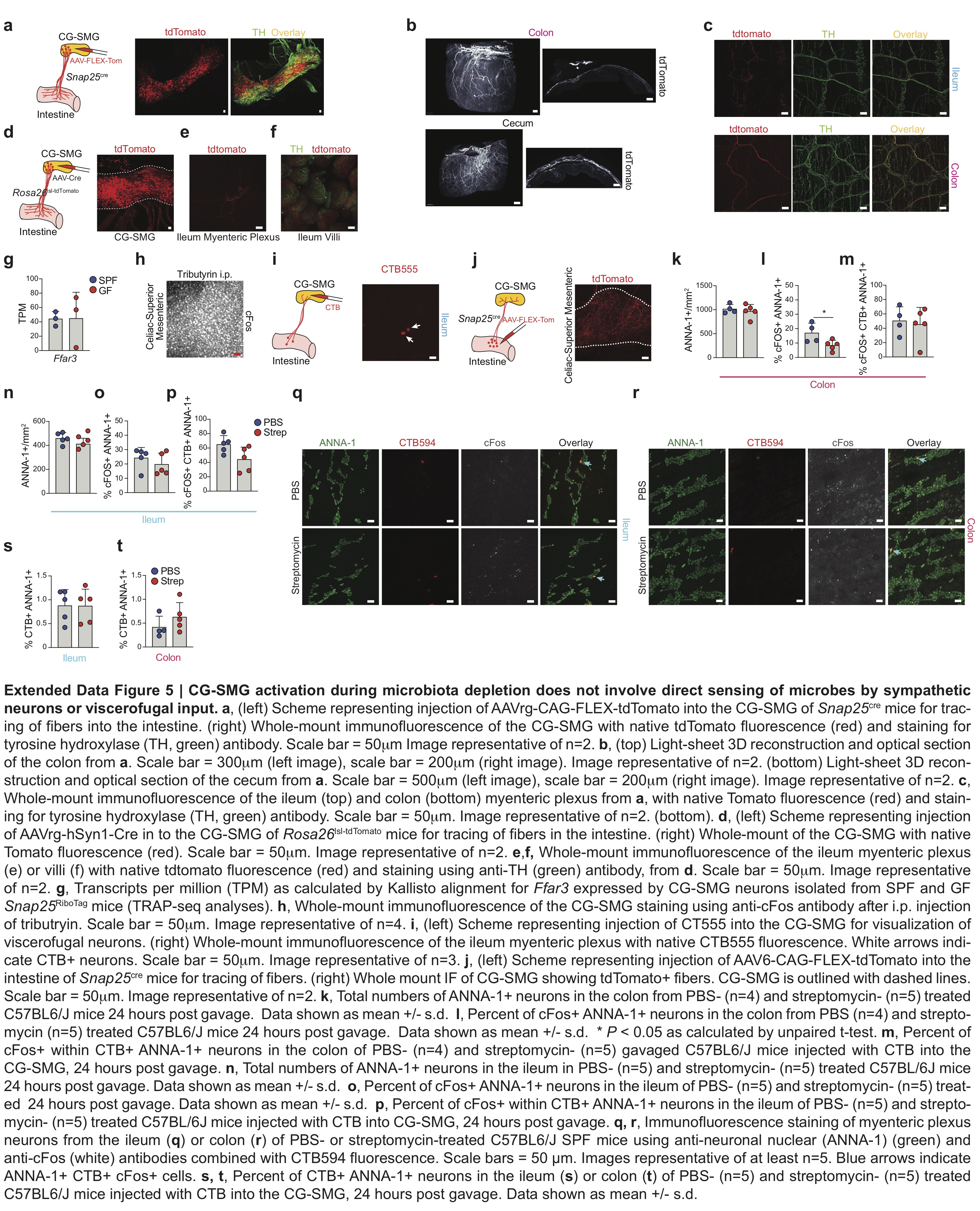

### Extended Fig6

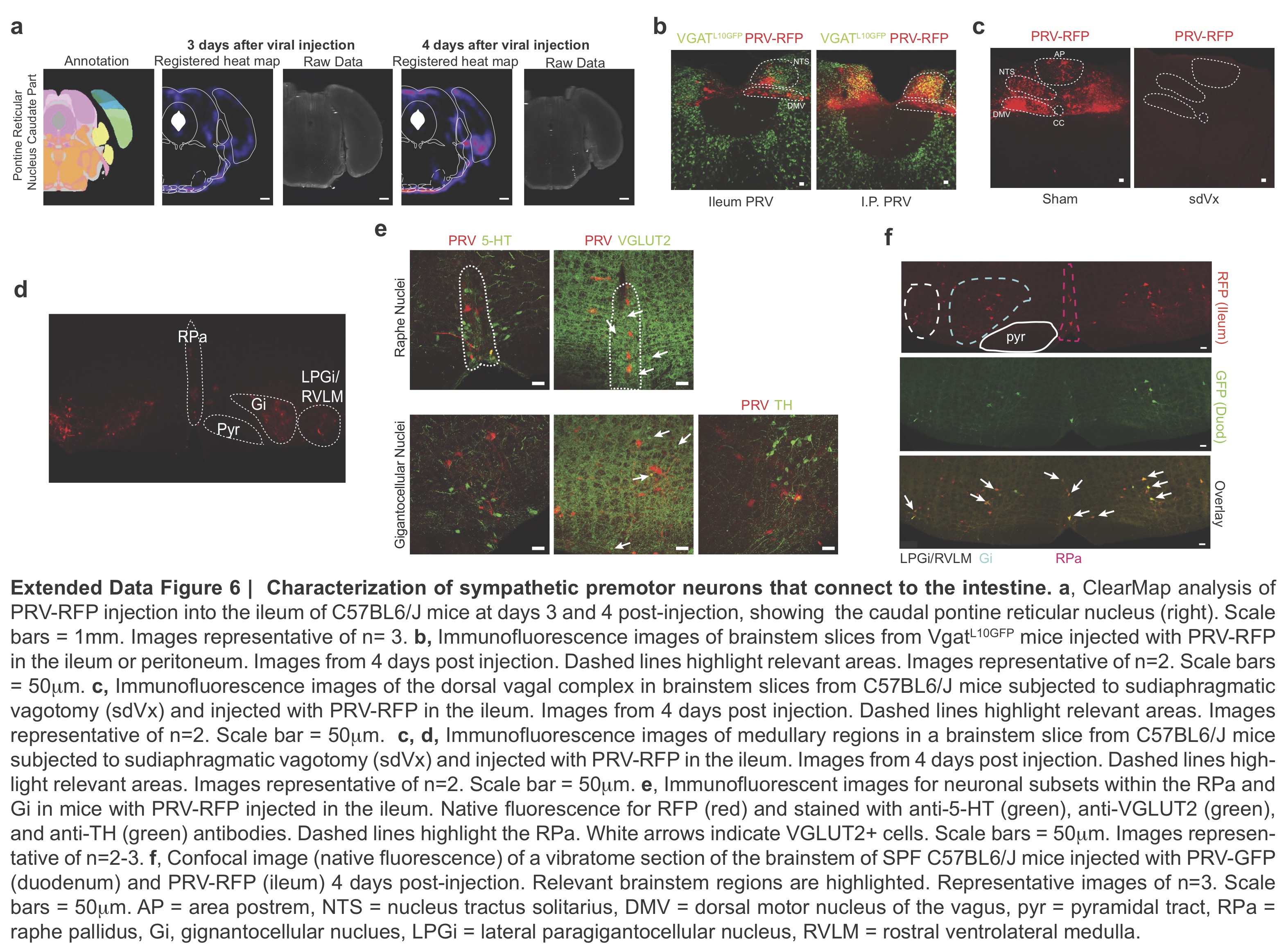

### Extended Fig7

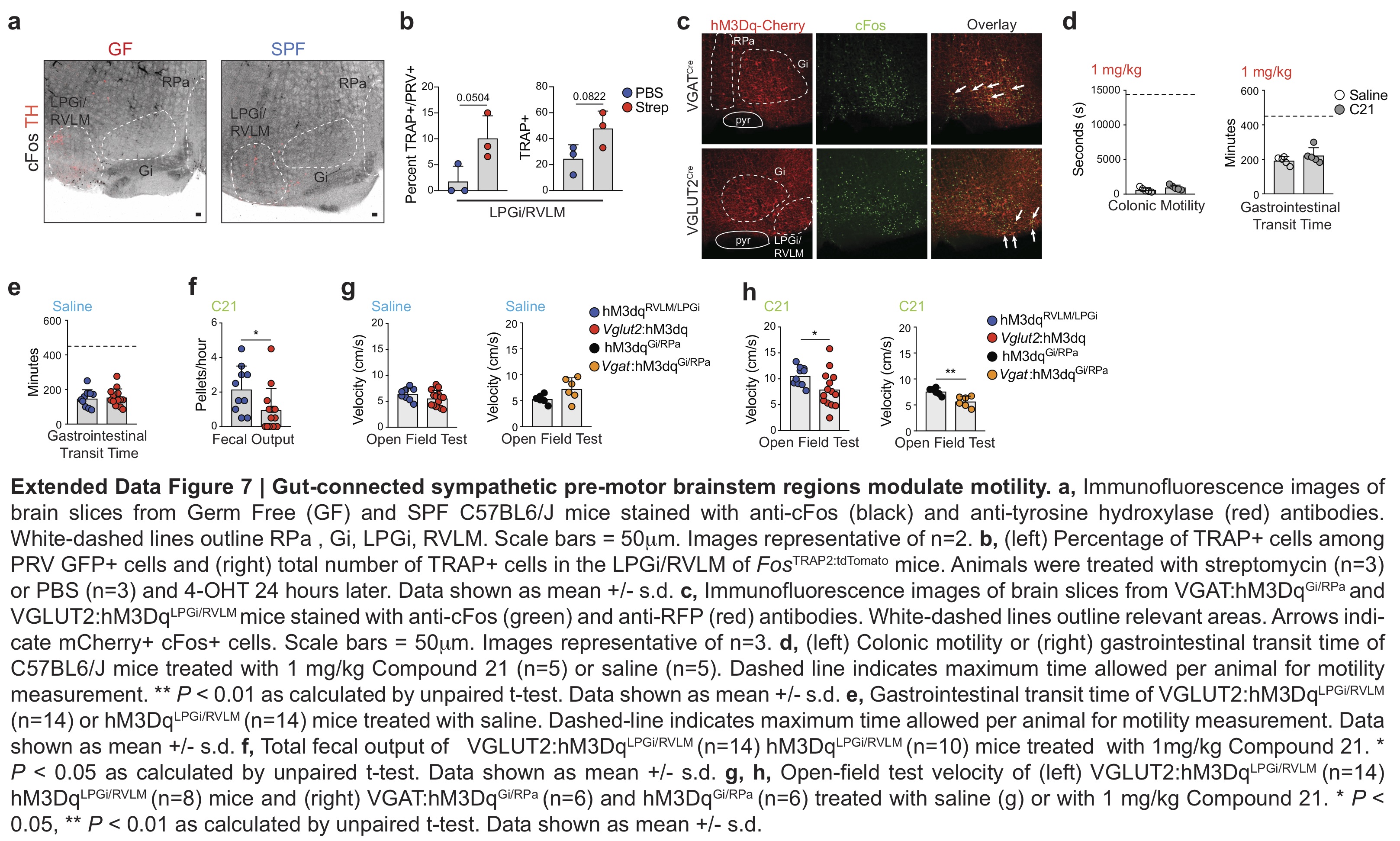

### Extended Fig8

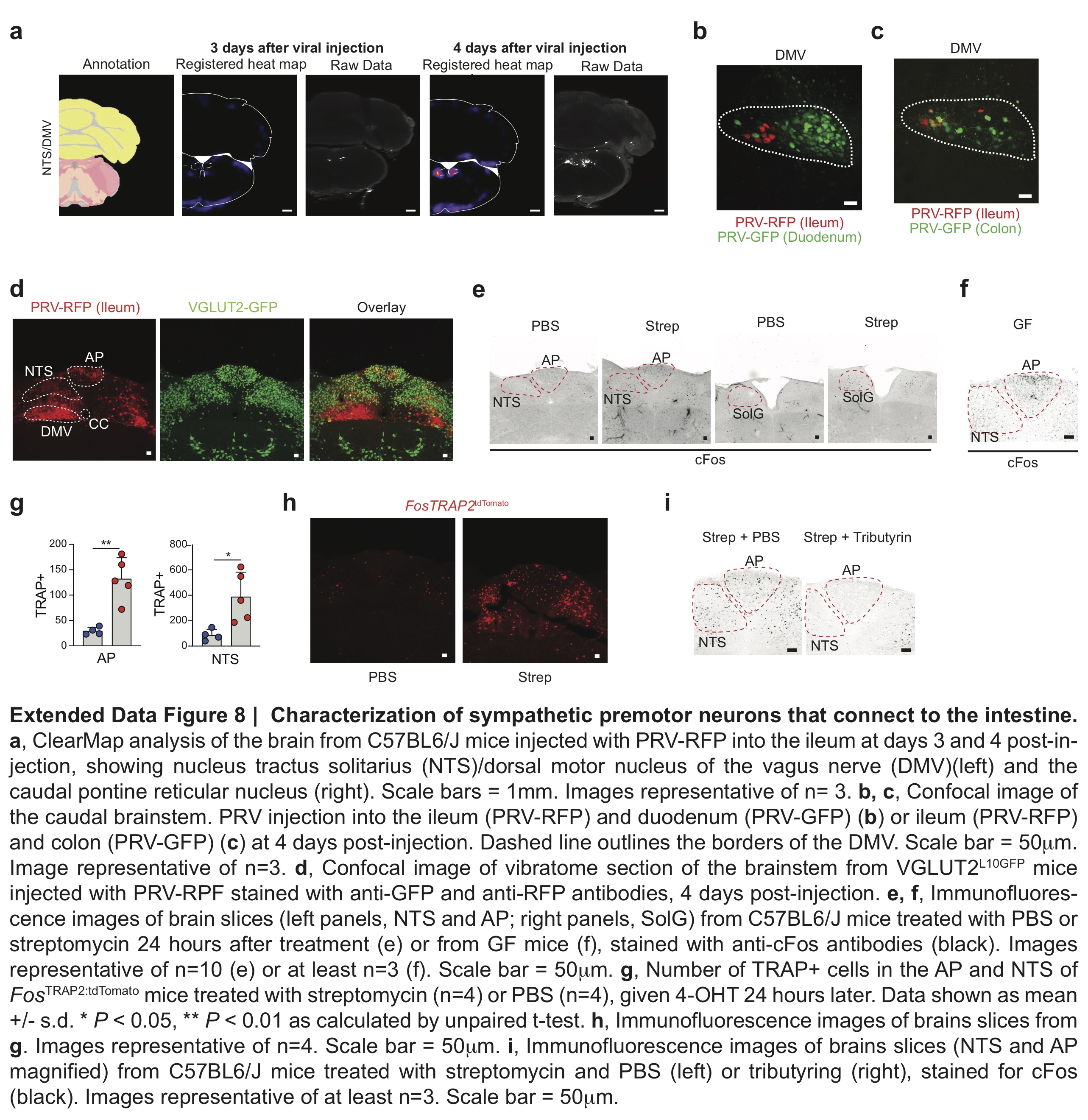

### Extended Fig9

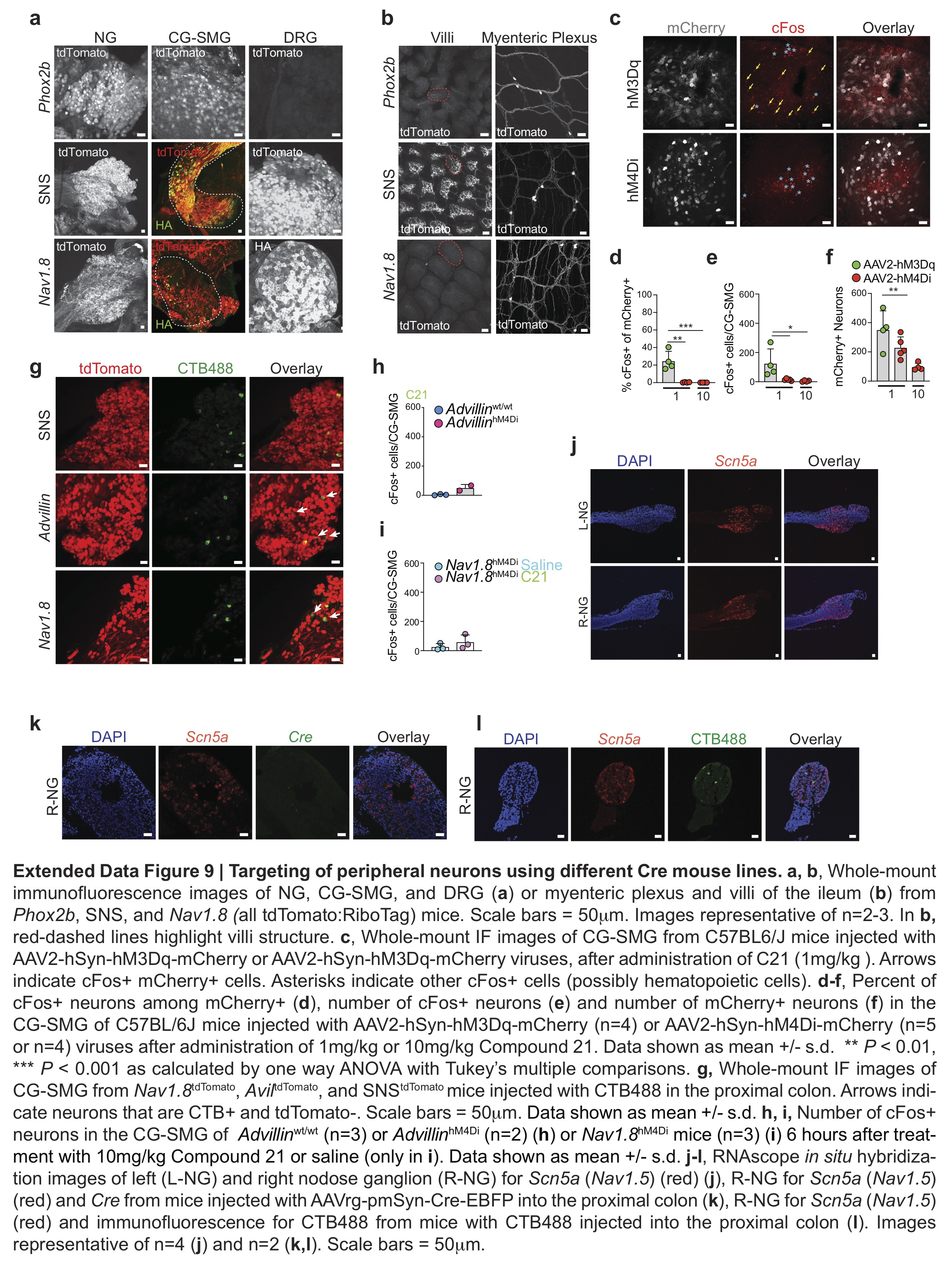

### Extended Fig10

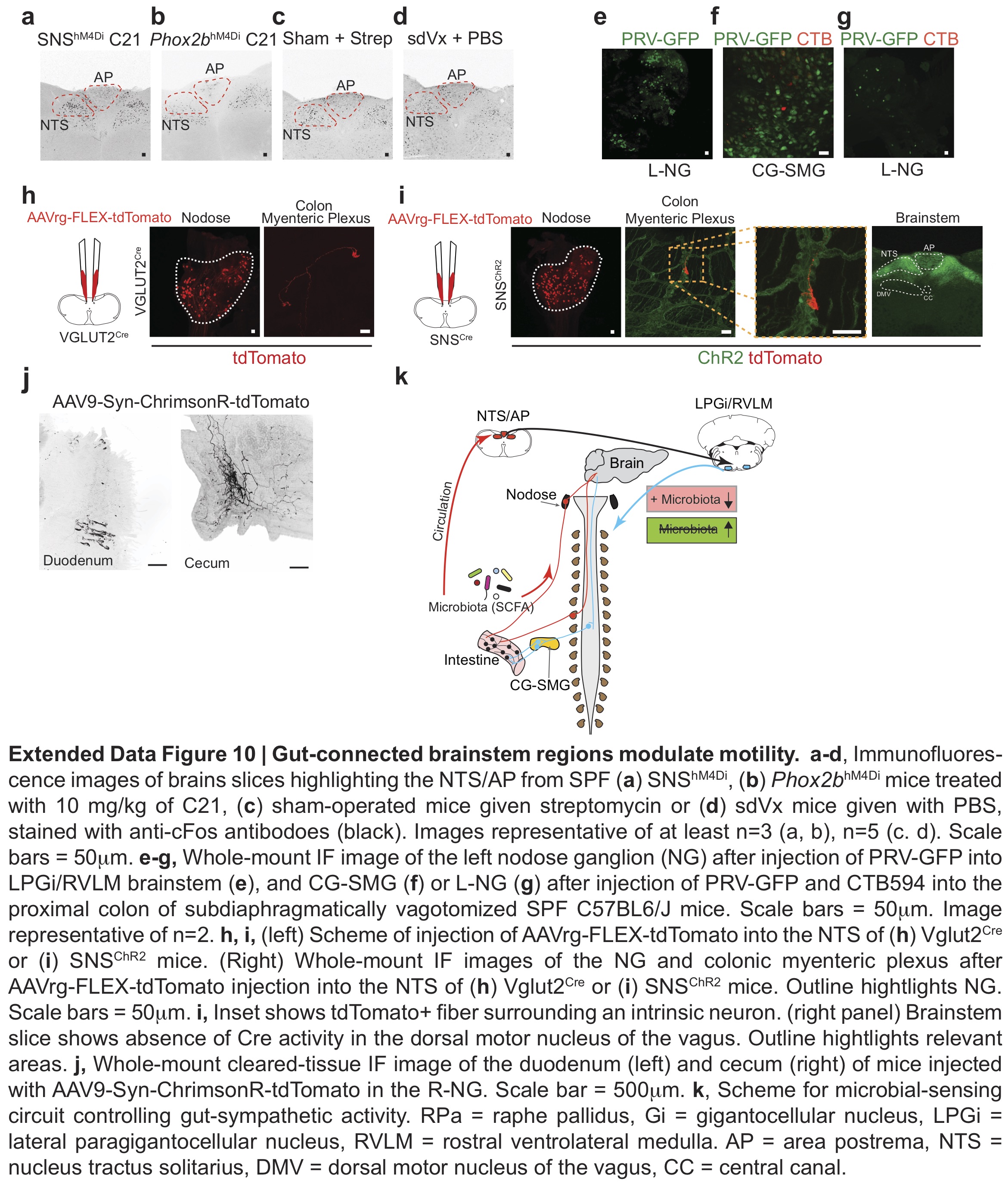

### Supplemental Annotations

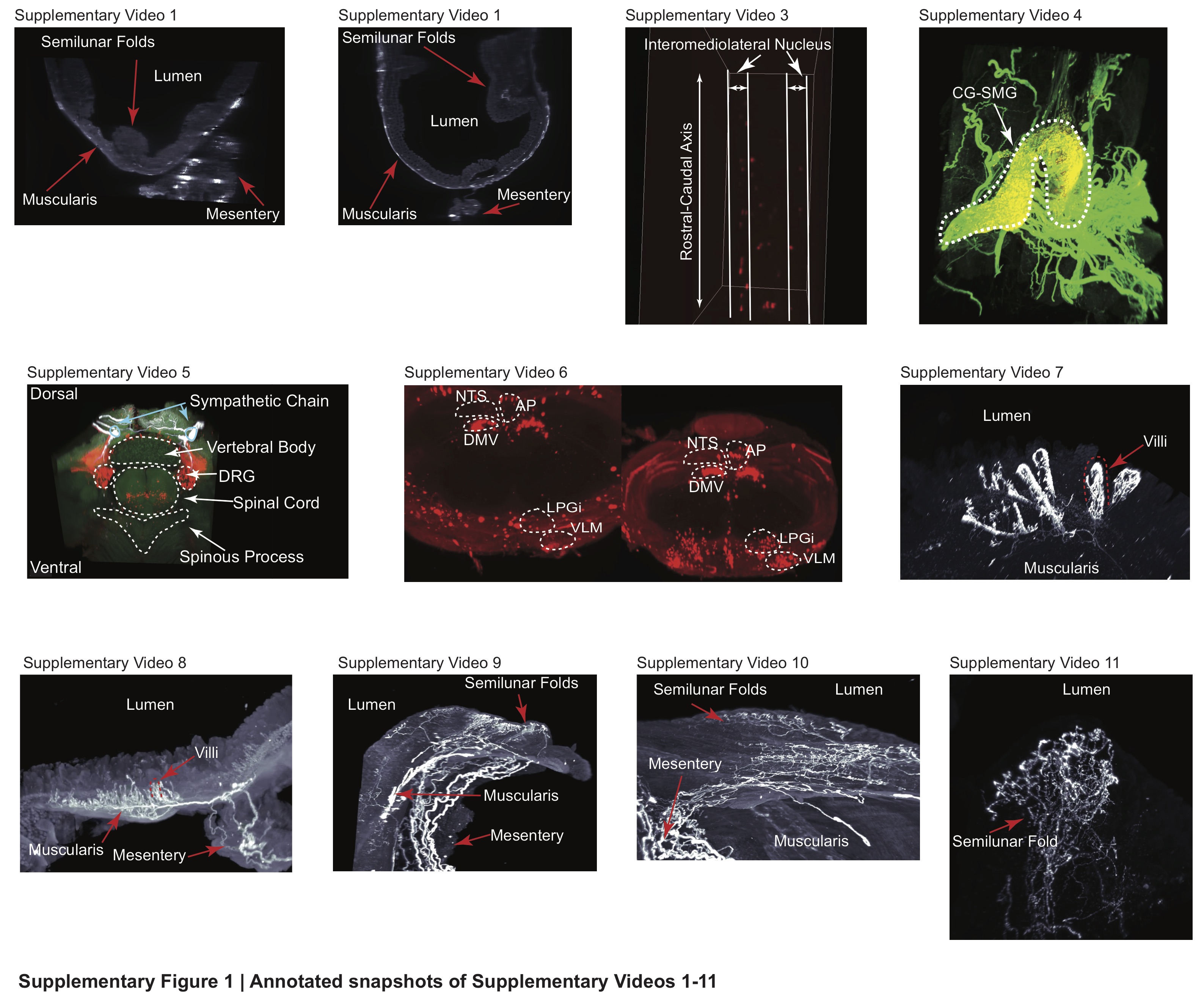

### Supplemental Table 1

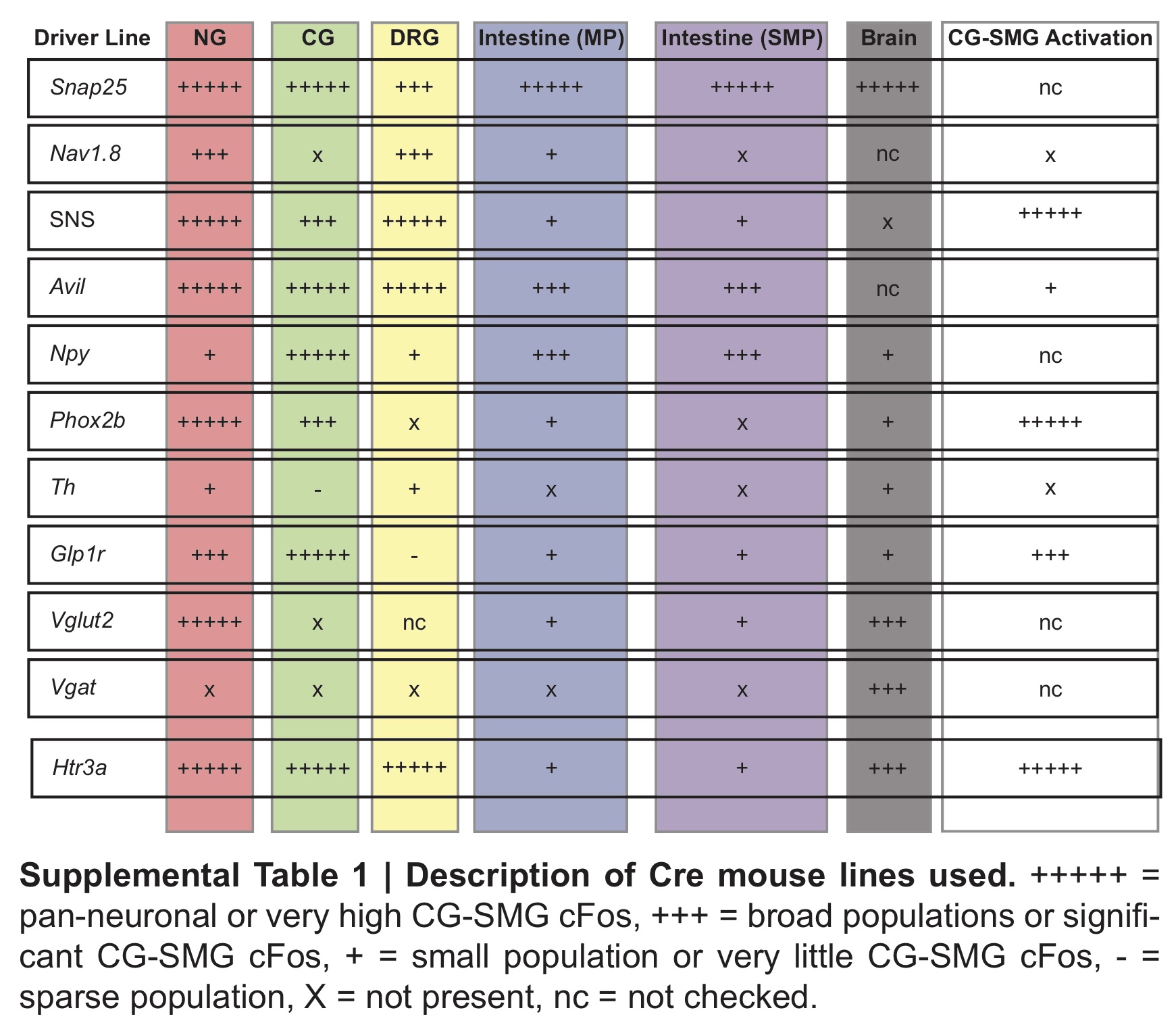
