## Supplemental Information 1 for "Microbes modulate sympathetic neurons via a gut-brain circuit"

**Supplementary Information 1**

Epithelial cell subsets, particularly enteroendocrine cells, are directly exposed to microbial signals and are capable of sensing and transmitting luminal input to EANs via synaptic transmission[1] and/or via secretion of neuropeptides and neurotransmitters[2]. Recent evidence pointed to a crucial role for the microbiota in modulating three major pathways: upregulation of serotonin[3], downregulation of L cell–derived glucagon-like peptide 1 (GLP-1)[4], and up or downregulation of peptide YY (PYY)[5,6]. Because secretion of these molecules is particularly enriched in the distal small intestine and colon, locations also enriched for SCFA production, we examined their role in the microbial modulation of sympathetic neurons. We addressed a possible suppressive role for epithelial–derived serotonin by crossing *Tph1*$^{fl/fl}$ mice with mice expressing inducible Cre$^{ER}$ under the villin promoter (Villin$^{Tph1}$). Conditional depletion of the key enzyme for serotonin production in gut epithelial cells upon tamoxifen administration did not result in changes in cFos+ neurons in the CG-SMG (Extended Data Fig. 4ac-c). In contrast, administration of the GLP-1R agonist Exendin-4 increased cFos+ neurons in the CG-SMG, as well as the total gastrointestinal transit time (Extended Data Fig. 4d-f). To assess whether this pathway was required for microbial regulation of gut sympathetic neurons, we treated *Glp1r*$^{-/-}$ mice with broad-spectrum antibiotics or Splenda. We observed similar numbers of cFos+ neurons in the CG-SMG isolated from the antibiotic-treated group as compared to wild-type mice, suggesting that GLP-1 is sufficient to drive sympathetic activity, albeit not required for microbial–dependent modulation of gut sympathetic neurons (Extended Data Fig. 4g). Finally, PYY administration did not activate the CG-SMG in SPF animals, but efficiently prevented the increase in cFos+ CG-SMG neurons following treatment with streptomycin (Extended Data Fig. 4g,i). These data suggest that the neuropeptides GLP-1 and PYY can modulate activity of gut sympathetic neurons and, in the case of PYY, may contribute to microbial regulation of gut sympathetic activity.
