## Supplemental Information 2 for "Microbes modulate sympathetic neurons via a gut-brain circuit"

**Supplementary Information 2**

We evaluated three possible neuronal inputs that could result in microbiota–dependent modulation of gut sympathetic neurons: i) direct sensing of the intestinal environment by CG-SMG sympathetic neurons due to innervation of the GI tract[1,2] and expression of Gpr41 by sympathetic neurons[3]; ii) changes in iEAN activity that could influence sensory afferents or sympathetic neurons directly (viscerofugal neurons)[4-6]; iii) sensory afferent modulation via the CNS.

i) To visualize the extent to which sympathetic innervation reaches the gut epithelium, we injected retrograde adeno-associated virus AAVrg-CAG-FLEX-tdTomato into the CG-SMG of *Snap25*$^{Cre}$ mice, thereby restricting the expression of Tomato protein to neurons (Extended Data Fig. 5a). As an independent and complementary strategy, AAVrg-hSyn1-Cre was injected into the CG-SMG of *Rosa*$^{lsl\text{-}tdTomato}$ mice. In both approaches, we observed restriction of viruses to the ganglion and the immediate surrounding tissue (Extended Data Fig 5b). Whole mount immunofluorescence or cleared–tissue imaging of the distal intestine showed innervation of the two neuronal plexuses, with sparse fibers extending to the mucosa, but no evidence of direct epithelial contact (Extended Data Fig. 5c-f, Supplementary Video 1,2). These results indicate that sympathetic eEANs may not be positioned to directly detect signals from the epithelial layer. It remains plausible that signals such as butyrate diffuse through the gut tissue or act via the circulation to influence CG-SMG activity. However, systemic (intraperitoneal) administration of tributyrin instead led to robust activation of the CG-SMG (Extended Data Fig. 5g), as previously suggested by expression of Gpr41 in sympathetic neurons[4] and confirmed by our TRAP-seq data (Extended Data Fig. 5h)[3]. Coupled with *Gpr41*$^{-/-}$ data, which showed CG-SMG activation, the above data *suggest* that direct detection by sympathetic neurons is not the primary mechanism of microbiota–mediated suppression of CG-SMG activation.

ii) Viscerofugal neurons are thought to be activated by tissue distension and could provide excitatory or inhibitory inputs to CG-SMG neurons[7,8]. To address whether viscerofugal neurons directly regulate microbiota–dependent CG-SMG activation, we injected fluorescent CTB into the CG-SMG and identified a sparse population of CTB$^+$ intrinsic EANs (iEANs), likely viscerofugal[7], in the intestine muscularis[7] (Extended Data Fig. 5i). To confirm that iEANs can make synaptic contacts with sympathetic neurons, we injected iEAN-tropic AAV6-CAG-FLEX-tdTomato[9] into the ileum of *Snap25*$^{cre}$ mice (Extended Data Fig. 5j). We detected tdTomato+ fibres within the CG-SMG, suggesting that viscerofugal neurons were capable of direct communication with sympathetic neurons (Extended Data Fig. 5j). To assess a possible link between activation of iEANs and the CG-SMG, we quantified the activation state of iEANs and viscerofugal neurons upon antibiotic treatment using cFos expression[10,11]. We injected fluorescent CTB into the CG-SMG of wild-type mice to retrograde label viscerofugal neurons. Oral gavage of streptomycin did not result in a change in the total number of neurons or viscerofugal neurons, or in an increase of cFos+ viscerofugal neurons in the distal intestine (Extended Data Fig. 5k-t). We concluded that viscerofugal iEANs are likely not playing a direct role in modulation of CG-SMG neurons; however, because changes in the microbial load can rapidly affect the activation state of iEANs, it remains possible that iEANs could influence sensory neuronal pathways[12].
