## Supplemental Information 3 for "Microbes modulate sympathetic neurons via a gut-brain circuit"

**Supplementary Information 3**

We found that both $SNS^{Cre}$ and $\mathit{Phox2b}^{Cre}$ also targeted a subset of sympathetic neurons in the CG-SMG, raising the possibility that the effects observed in both mice could be due to direct effects in these ganglia (Extended Data Fig 9a, b, Supplemental Table 1). This is an unlikely possibility, since sympathetic neurons in the CG-SMG do not noticeably express either *Girk2*, a key component of hM4Di-driven inhibition[1], or *Chrm4*, the endogenous cholinergic muscarinic receptor (*data not shown*). Nevertheless, to address this issue, we asked whether expression of excitatory hM3Dq or inhibitory hM4Di specifically in CG-SMG neurons could result in cFos expression following C21 administration. We injected AAV2-hSyn-hM4Di or hM3Dq into the CG-SMG of wild-type mice and administered either 1mg/kg or 10mg/kg of C21. We observed robust increase in cFos expression by hM3Dq+ sympathetic neurons, but we failed to detect cFos+ hM4Di+ with either dose of C21 (Extended Data Fig. 9c-f). Extensive analyses of additional Cre lines revealed different patterns of NG targeting and CG-SMG activation, particularly with $\mathit{Nav1.8}^{Cre}$, which targets Scn10a, and with the pan-sensory $\mathit{Advillin}^{CreERT2}$ (Supplemental Table 1).
